# Automated continuous evolution of proteins *in vivo*

**DOI:** 10.1101/2020.02.21.960328

**Authors:** Ziwei Zhong, Brandon G. Wong, Arjun Ravikumar, Garri A. Arzumanyan, Ahmad S. Khalil, Chang C. Liu

## Abstract

We present automated continuous evolution (ACE), a platform for the hands-free directed evolution of biomolecules. ACE pairs OrthoRep, a genetic system for continuous targeted mutagenesis of user-selected genes *in vivo*, with eVOLVER, a scalable and automated continuous culture device for precise, multi-parameter regulation of growth conditions. By implementing real-time feedback-controlled tuning of selection stringency with eVOLVER, genes of interest encoded on OrthoRep autonomously traversed multi-mutation adaptive pathways to reach desired functions, including drug resistance and improved enzyme activity. The durability, scalability, and speed of biomolecular evolution with ACE should be broadly applicable to protein engineering as well as prospective studies on how selection parameters and schedules shape adaptation.

---

Continuous evolution has emerged as a powerful paradigm for the evolution of proteins and enzymes^1–3^ towards challenging functions^4,5^. In contrast to classical directed evolution approaches that rely on stepwise rounds of *ex vivo* mutagenesis, transformation into cells, and selection^6^, continuous evolution systems achieve rapid diversification and functional selection autonomously, often through *in vivo* targeted mutagenesis systems (Fig. 1a). The result is a mode of directed evolution that requires only the basic culturing of cells, in theory, enabling extensive speed, scale, and depth in evolutionary search^3^. In practice, however, developing a continuous evolution method that realizes all three properties has been challenging.

**Figure 1.**
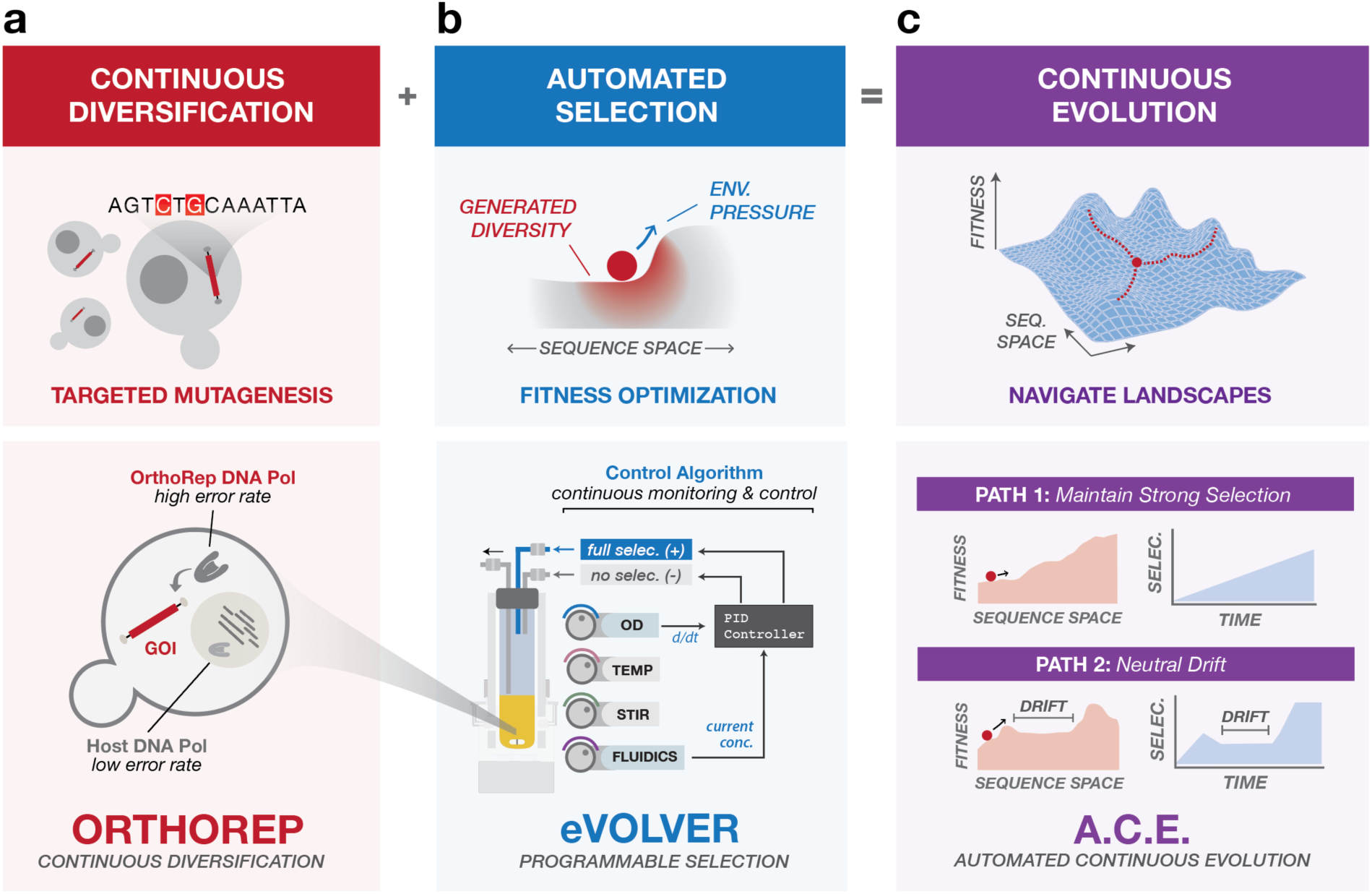
Automated Continuous Evolution (ACE). (**a**) OrthoRep enables continuous diversification of genes of interest (GOIs) via *in vivo* targeted mutagenesis in yeast. The basis of OrthoRep is an orthogonal DNA polymerase-plasmid pair that mutates GOIs ~100,000-fold faster than the genome. (**b**) eVOLVER is a continuous culturing platform for programmable, multiparameter control of selection conditions across many independent cultures. A PID control algorithm implemented with eVOLVER dynamically tunes selection pressure of populations as they adapt, precisely challenging them to achieve desired functions. PID control is achieved by tuning the ratio of full selection and no selection media inputs in response to growth rate. (**c**) By running OrthoRep in eVOLVER with PID control, ACE autonomously and rapidly navigates complex fitness landscapes. With a single framework, ACE can guide independent cultures through diverse trajectories.

Recently, our groups made two independent advances that can pair to achieve continuous evolution at significant speed, scale, and depth. These advances are OrthoRep and eVOLVER. First, OrthoRep. OrthoRep is an engineered genetic system for continuous *in vivo* targeted mutagenesis of genes of interest (GOIs)^2,7^. OrthoRep uses a highly error-prone, orthogonal DNA polymerase-plasmid pair in yeast that replicates GOIs at a mutation rate of 10^−5^ substitutions per base (spb) without increasing the genomic mutation rate of 10^−10^ spb (Fig. 1a). This ~100,000-fold increase in the mutation rate of GOIs drives their accelerated evolution (speed). Because the OrthoRep system functions entirely *in vivo* and culturing yeast is straightforward, independent GOI evolution experiments can be carried out in high-throughput (scale). In addition, long multi-mutation pathways can be traversed using OrthoRep, owing to the durability of mutagenesis over many generations (depth). However, to practically realize depth in evolutionary search, *in vivo* mutagenesis with OrthoRep must be coupled with a functional selection that can be tuned over the course of a continuous evolution experiment. This tuning is necessary to precisely and efficiently guide populations to the desired evolutionary search depth. For example, evolution of novel functions requiring long mutational trajectories may demand frequent modification of selection conditions in order to maintain strong selection^4,5,8^, guide evolution through strategic intermediate functions^1,5^, or impose periods of neutral drift or alternating selection to promote crossing of fitness valleys^9,10^ (Fig. 1c). Yet, selection schedules cannot be determined *a priori* as the generation of beneficial mutations is a fundamentally stochastic process. Therefore, selection schedules should be adjusted dynamically based on how populations adapt, rendering manual implementation of continuous evolution experiments onerous. Previous continuous evolution campaigns approached the challenge of optimizing selection schedules by either limiting the number of parallel evolution experiments being carried out so that selection can be manually changed on the fly^1,4^, or by setting a fixed but conservative selection schedule to buffer against variations in adaptation rate across a large numbers of replicate experiments^2^. However, even with conservative selection schedules, a proportion of replicates in high-throughput evolution studies went extinct when the rate of selection stringency increase outpaced the rate of adaption^2^. Enter eVOLVER. eVOLVER is a versatile continuous culture platform that enables multiparameter control of growth selection conditions across independent microbial cultures^11,12^ (Fig. 1b). eVOLVER’s flexible hardware and software permit development of “algorithmic selection routines” that apply selective pressures based on real-time monitoring and feedback from culture growth characteristics. Additionally, eVOLVER’s robust framework ensures experimental durability over long timeframes, and its unique scalable design allows independent control over tens to hundreds of cultures. Combining OrthoRep and eVOLVER should therefore enable continuous evolution with speed, depth, and scale.

Here we describe this pairing of OrthoRep with eVOLVER to achieve <u>A</u>utomated <u>C</u>ontinuous <u>E</u>volution (ACE) (Fig. 1c). By implementing a closed-loop feedback routine that dynamically adjusts the strength of selection for a desired function in response to growth rate changes of yeast populations diversifying a GOI on OrthoRep, we demonstrate continuous evolution over extended periods of time without manual intervention. To illustrate the performance and utility of ACE, we describe its application in two model protein evolution experiments, one yielding drug-resistant *Plasmodium falciparum* dihydrofolate reductases (*Pf*DHFRs) and the other yielding variants of the thermophilic HisA enzyme from *Thermotoga maritima* (*Tm*HisA) that operate well in mesophilic yeast hosts.

To establish ACE, we first reconfigured eVOLVER Smart Sleeves^13^ so that each culture vial receives two media inputs: (1) ‘no selection’ base media (*e.g.* media without drug or with the maximum concentration of nutrient in our cases) and (2) ‘full selection’ media (*e.g.* media with the maximum concentration of drug or without nutrient in our cases). As a result, using eVOLVER software calculations, selection strength can be dynamically tuned by altering the ratios of the two media inputs as cultures are diluted over time (Fig. 1b,c). We then implemented a closed-loop control system that seeks to achieve and maintain a target culture growth rate by dynamically adjusting selection strength. Briefly, culture growth rate is continuously measured based on real-time recordings of optical density (OD), and a proportional-integral-derivative (PID) control algorithm^13^ is used to determine the percentage of full selection media to add to the culture in order to minimize error between the actual growth rate and a target growth rate (or setpoint) (see **Methods**). Although simpler feedback algorithms have been previously used in microbial evolution experiments^14,15^, these resulted in growth rate oscillations or excessive overshooting in our experiments, frequently driving cultures to extinction (Supplementary Fig. 1).

To validate ACE, we first repeated a continuous evolution experiment that we previously conducted using manual serial passaging. Specifically, we evolved *Plasmodium falciparum* dihydrofolate reductase (*Pf*DHFR) to develop drug resistance to the antimalarial drug, pyrimethamine, by encoding *Pf*DHFR on OrthoRep in a yeast strain that relies on *Pf*DHFR activity for survival^2^ (Fig. 2). We determined appropriate PID constants to tune the concentration of pyrimethamine (Fig. 2b and Supplementary Fig. 2) and keep the measured growth rate of cells at a target growth rate (Fig. 2a, setpoint = dashed black line). This program forced cells to continuously experience a strong selection pressure imposed by pyrimethamine, which resulted in the rapid evolution of *Pf*DHFR resistance (Fig. 2c). We observed that after ~550 hours (~100 generations) of continuous hands-free operation of ACE, five out of six replicates adapted to 3 mM pyrimethamine, the highest concentration of pyrimethamine soluble in liquid media (Fig. 2b and Supplementary Fig. 3). ACE maintained cultures near the target growth rate over the entire course of the experiment (Fig. 2a,b), demonstrating the effectiveness of the control loop. In contrast to the use of a fixed selection schedule^2^ or simpler control algorithms for selection (Supplementary Fig. 1) that resulted in occasional extinction caused by too-rapid increases in pyrimethamine concentration, all six ACE experiments reliably adapted to yield multi-mutation pyrimethamine-resistant *Pf*DHFR variants. Validating our method, we found that populations converged on strong resistance mutations in *Pf*DHFR – C50R, D54N, Y57H, C59R, C59Y, and S108N – as observed and characterized previously^2^ (Fig. 2c). Additionally, the monotonically increasing pyrimethamine concentrations we observed for most replicates (Fig. 2b) are consistent with step-wise fixation of beneficial mutations expected for the evolution of *Pf*DHFR resistance under strong selection^2,16^. Finally, ACE demonstrated a substantial increase in speed over our previous evolution campaign performed by manual passaging; with ACE, culture growth rates in 5/6 vials stabilized at the maximum pyrimethamine concentration after ~550 hours, which is over 200 hours faster than for the manual evolution campaign done with serial passaging^2^. Collectively, these results validate the ACE system and highlight its ability to enable reliable and rapid continuous evolution of proteins.

**Figure 2.**
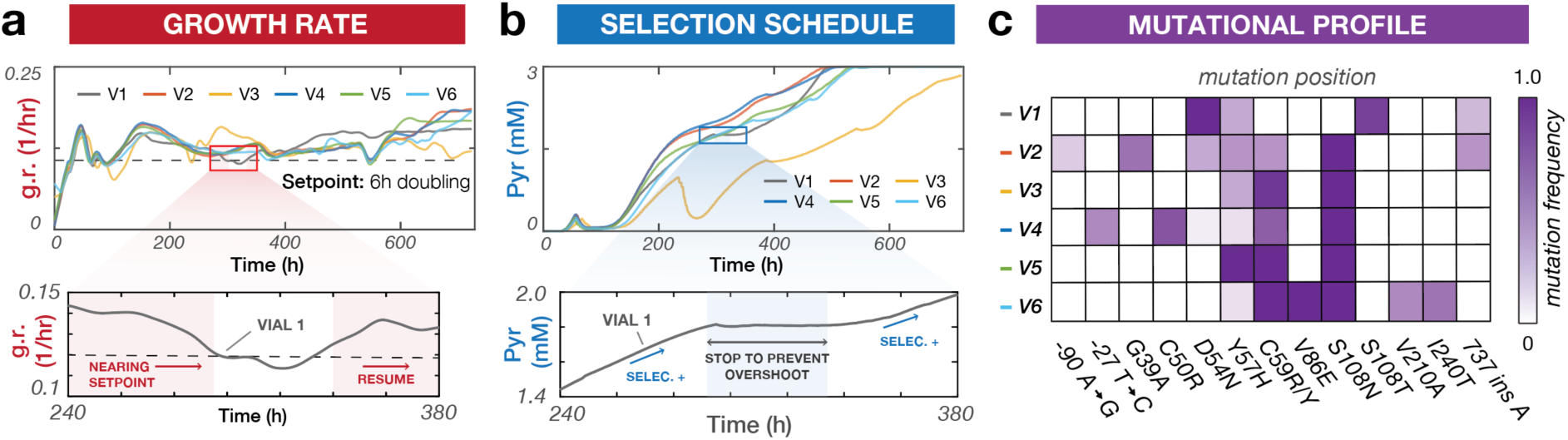
Automated continuous evolution of *Pf*DHFR resistance to pyrimethamine. (**a**) Top: Growth rate traces for six independent OrthoRep cultures (V1-6) evolving *Pf*DHFR resistance to pyrimethamine in eVOLVER using PID control. Bottom: A representative time window validating PID control. The growth rate (solid line) is controlled by automated tuning of pyrimethamine concentration (Fig. 2b, bottom) to keep cultures constantly challenged at the setpoint growth rate (dashed line). (**b**) Top: Drug selection schedules for OrthoRep cultures evolving *Pf*DHFR. Bottom: A representative time window demonstrating PID-based selection tuning. Pyrimethamine concentration autonomously adjusts in response to growth rate deviation from the setpoint (Fig. 2a, bottom). (**c**) Promoter and *Pf*DHFR mutations identified in six evolved populations. Mutation frequencies are estimated from SNP analysis of bulk Sanger sequencing traces.

**Figure 3.**
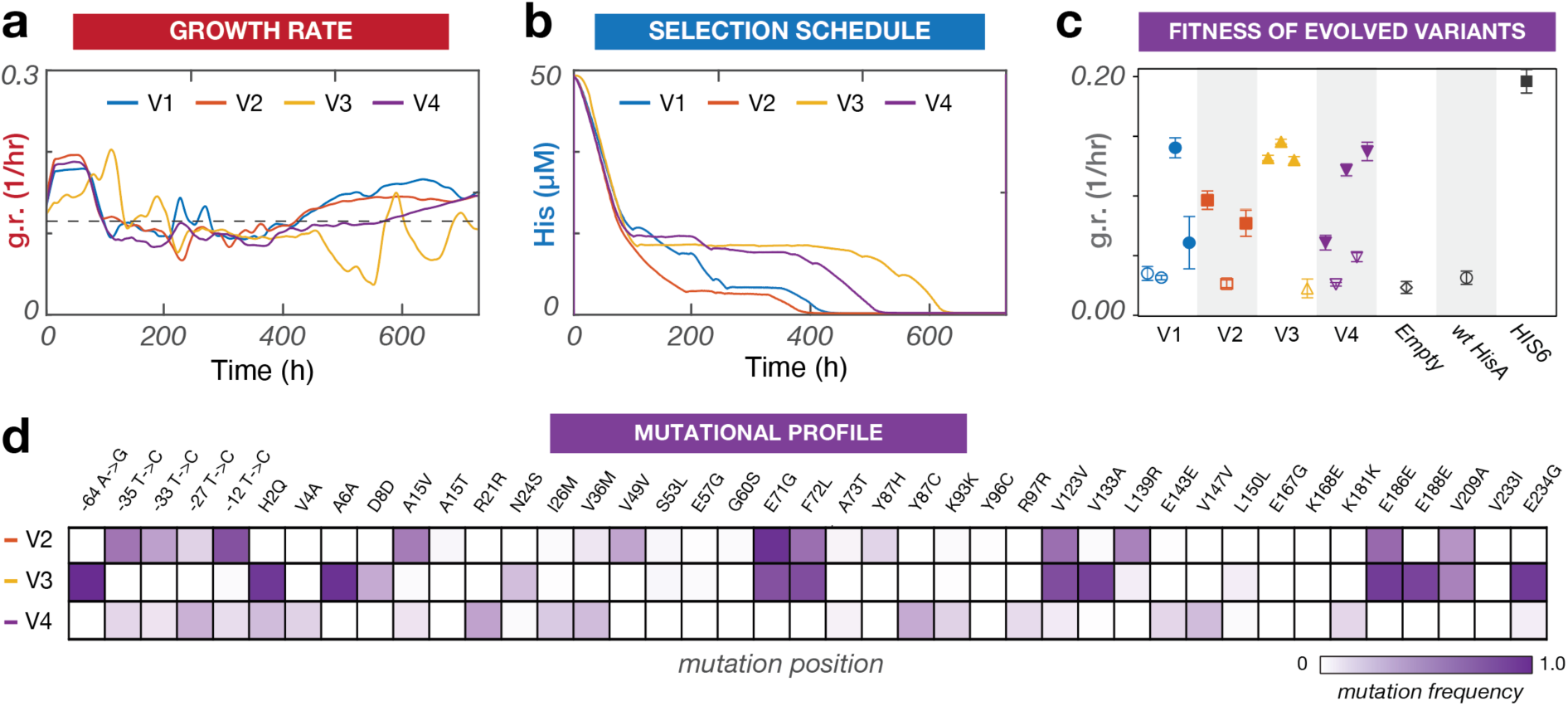
Automated continuous evolution of *Tm*HisA to operate in mesophilic yeast. (**a**) Growth rate traces of four independent OrthoRep cultures (V1-4) evolving *Tm*HisA to support growth of a Δ*his6* yeast strain at 30 **°**C. (**b**) Nutrient concentration schedule of the four evolving cultures controlled via PID. (**c**) Growth rate analysis of individual *Tm*HisA variants selected from adapted cultures. Three to five *Tm*HisA variants were sampled from each replicate evolution experiment. The coding regions of the variants were cloned into a low-copy yeast plasmid under the control of a medium strength promoter. The ability of the variants to support growth of a Δ*his6* strain was measured in triplicate in comparison to the ability of wt *Tm*HisA and native yeast HIS6 to support growth. Shapes indicate means and error bars denote standard deviations. Darkened replicates indicate a p < 0.05 compared to wt *Tm*HisA activity by one-way ANOVA. We note that a number of evolved variants sampled from our adapted replicates did not support growth, but this is likely because OrthoRep evolves *Tm*HisA in the context of a multi-copy orthogonal plasmid, allowing inactive variants to hitchhike with active copies in the same cell. Such inactive variants could be sampled during the subcloning of *Tm*HisA from orthogonal plasmids into the low-copy plasmids used for testing variants in fresh strains. (**d**) Promoter and *Tm*HisA mutations identified in evolved populations from V2-4. Mutation frequencies are estimated from SNP analysis of bulk Sanger sequencing traces. V1 Sanger sequencing traces are not included due to a technical mistake that rendered stocks of the evolved population inviable for revival and sequencing, although individual clones were fully analyzed (Fig. 3c), as they were sampled before the stocking mistake.

We next applied ACE to evolve the thermophilic *Thermotoga maritima* HisA enzyme (*Tm*HisA) to function in *Saccharomyces cerevisiae* at mesophilic temperatures. *Tm*HisA, an ortholog of *S. cerevisiae* HIS6, catalyzes the isomerization of ProFAR to PRFAR in the biosynthesis of histidine. However, wild-type *Tm*HisA does not effectively complement a *his6* deletion in yeast when expressed from a medium-strength yeast promoter (Fig. 3), likely due to the different temperature niches of *S. cerevisiae* and *T. maritima* (30°C and 80°C, respectively). We reasoned that ACE could readily drive the evolution of *Tm*HisA to function in yeast Δ*his6* strains by selecting for growth in media lacking histidine. This evolution serves as a valuable test of the capabilities of ACE for at least two reasons. First, adapting enzymes from non-model thermophiles to function in model mesophiles is useful for industrial biotechnology whose infrastructure is designed around model organisms like yeast and bacteria. Second, in contrast to drug resistance in *Pf*DHFR, which is driven by a small number of large effect mutations^3^, we reasoned that temperature adaptation of enzyme activity would involve a large number of small effect mutations, leading to a more complex fitness landscape. This would act as a demanding test of ACE requiring precise feedback-control to efficiently traverse adaptive paths.

We encoded *Tm*HisA on OrthoRep in a Δ*his6* strain and carried out ACE selection in four independent replicates for a total of 600 hours (~100 generations) (Fig. 3 and Supplementary Fig. 4). At the beginning of the experiment, there was no detectable growth in the absence of histidine. At the end of the experiment, all four replicates successfully adapted to media lacking histidine. To confirm that *Tm*HisA evolution was responsible for the observed adaptation, *Tm*HisA variants were isolated from OrthoRep and characterized for their ability to complement a *his6* deletion in fresh yeast strains. Indeed, the evolved *Tm*HisA variants we sampled were able to support growth in media lacking histidine in contrast to wild-type *Tm*HisA (Fig. 3c). Consistent with a model of a more complex fitness landscape, growth rate traces for the four replicate cultures were noisier (Fig. 3a) than those of *Pf*DHFR (Fig. 2a), full adaptation occurred only after a long period of neutral drift (hours ~100-500 in Fig. 3b), and the sequences of independently evolved *Tm*HisAs were diverse (Fig. 3d and Supplementary Table 1). Nevertheless, ACE was able to autonomously adapt *Tm*HisA in all four replicates within 120 fewer hours than manual passaging experiments (unpublished results). Sequencing of sampled clones revealed *Tm*HisA variants harboring between 6 and 15 mutations (Supplementary Table 1), again demonstrating the durability of ACE in carrying out long evolutionary searches to discover high-activity multi-mutation enzyme variants.

In summary, we have developed a fully automated, *in vivo* continuous evolution setup termed ACE that couples OrthoRep-driven continuous mutagenesis and eVOLVER-enabled programmable selection. We demonstrated the evolution of drug resistance in *Pf*DHFR and mesophilic operation of *Tm*HisA, showcasing the ability of ACE to individually control selection schedules in multi-replicate GOI evolution experiments based on real-time measures of adaptation. The result is a system that offers unprecedented speed, depth, and scalability for conducting evolutionary campaigns to achieve ambitious protein functions.

ACE paves the way for an array of complex evolution experiments that can advance both basic and applied protein and enzyme evolution. For example, eVOLVER can be used to program multidimensional selection gradients across OrthoRep experiments, test the effects of different population sizes (and beneficial mutation supply) on the outcomes of adaptation, or explore the relationship between timescales of drift and adaptation. Real-time feedback on growth metrics to adjust selection stringency can ensure that every evolving population is being constantly challenged or allowed to drift, which is especially relevant when evolving biomolecules with rugged fitness landscapes where predefined selection strategies are prone to driving populations to extinction or local fitness maxima. In the future, many other algorithmic selection routines may be implemented with ACE to more efficiently and intelligently navigate fitness landscapes. For example, machine learning algorithms can take the outcomes of replicate evolution experiments carried out under different selection schedules to train ACE selection programs themselves. Finally, the automated, open-source nature of ACE is well-suited for integration with other open-source hardware and wetware tools to create larger automation pipelines. Overall, we foresee ACE as an enabling platform for rapid, deep, and scalable continuous GOI evolution for applied protein engineering and studying the fundamentals of protein evolution.

## Supporting information

Code associated with main paper

## Data availability

The raw data that supports the plots and other findings within this paper are available in the supplementary information and from corresponding authors upon reasonable request.

## Code availability

The code for eVOLVER operation, including feedback control of selection stringency, is available as supplementary files in the online version of this manuscript. The MATLAB script for growth rate determination is available as a supplementary file in the online version of this manuscript.

## Acknowledgements

We thank members of the Liu and Khalil groups for helpful discussions. This work was funded by NIH (1DP2GM119163-01), NSF (MCB1545158), and DARPA (HR0011-15-2-0031) to CCL, and NIH (1DP2AI131083-01, 1R01EB027793-01), NSF (CCF-1522074), and DARPA (HR0011-15-C-0091) to ASK. We especially thank Dr. Justin Gallivan and the DARPA Biological Robustness in Complex Settings program, which sponsored the construction of an eVOLVER system in the Liu lab.

## Author contributions

All authors designed experiments. ZZ, BGW, AR, GAA, and CCL performed experiments. All authors analyzed results. ZZ and BGW programmed the eVOLVER system and wrote code. GAA wrote the MATLAB script for growth rate analysis. ZZ, BGW, AR, ASK, and CCL wrote the paper. ASK and CCL procured funding and oversaw the project.

## Supplementary Material Contents

Supplementary Figures

Supplementary Fig. 1. Sample evolution experiments in eVOLVER using the control algorithm derived from Toprak et al.

Supplementary Fig. 2. Oscillation of growth rate of ZZ-Y323 during the empirical determination of PID settings.

Supplementary Fig. 3. Adaptation history for all six replicates during *Pf*DHFR evolution.

Supplementary Fig. 4. Adaptation history for all four replicates during *Tm*HisA evolution.

Supplementary Tables

Supplementary Table 1. *Tm*HisA mutants characterized.

Supplementary Table 2. List of plasmids used in this study.

Supplementary Table 3. List of yeast strains used in this study.

## Methods

### Cloning

All plasmids used in this study are listed in Supplementary Table 2. Plasmids were cloned using either restriction enzymes if compatible sites were available or using Gibson cloning^17^ with 20-40 bps of overlap. Primers and gBlocks were ordered from IDT Technologies. Enzymes for PCR and cloning were purchased from NEB. Plasmids were cloned into either Top10 *E. coli* cells from Thermo Fisher or SS320 *E. coli* from Lucigen.

### Yeast transformation and DNA extraction

All yeast strains used in this study are listed in Supplementary Table 3. Yeast transformations were done with roughly 100 ng – 1 μg of plasmid or donor DNA via the Gietz high-efficiency transformation method^18^. For integration of genes onto the orthogonal plasmid (pGKL1), cassettes were linearized with ScaI and subsequently transformed as described previously^2,7^. Standard preparations of YPD and drop-out synthetic media were obtained from US Biological. When necessary, the following were supplemented at their respective concentrations: 5-FOA at 1 mg/mL, G418 at 400 µg/mL, and Nourseothricin at 200 µg/mL. Yeast DNA extraction of orthogonal plasmids were done as previously reported^2,7^.

### eVOLVER feedback control configuration

ACE experiments were performed using the previously described eVOLVER continuous culture system^11^, modified to enable an additional media input into each culture. Specifically, each vessel consists of three connected pumps (two input, one efflux) and are actuated programmatically to implement a so-called “morbidostat” algorithm where the selection stringency is adjusted to maintain a particular rate of cell growth. The custom script of eVOLVER (custom_script.py) was extensively modified to change the behavior of eVOLVER from the default turbidostat to a morbidostat. Briefly, in the new morbidostat mode, eVOLVER dilutes the growing cultures after a defined time, which we set to an hour. At the time of dilution, the growth rate since the last dilution is calculated by fitting the OD measurements to an exponential equation *y* = *A* ∙ *e*^*Bx*^ where *B* is the growth rate. Using the current and historical growth rate, a dilution parameter, *r*(*t*) was calculated as described below to dilute the morbidostat. The morbidostat algorithm and eVOLVER experimental code are written in Python and included in the supplemental files.

The efflux pump for each vessel is actuated whenever either of the influx pumps are triggered and stay ON for an additional 5 seconds. Therefore, the volume of the culture vessel is determined by the length of the efflux straw and estimated to be at 30 mL. The flow rate of each media input was individually calibrated for accurate metering of drug or nutrient into the culture.

Before each experiment, 40 mL borosilicate glass vessels (Chemglass), stir bars (Fisher), and fluidic straws were assembled and autoclaved. Fluidic lines were sterilized by flushing with 10% bleach and 70% ethanol before use. Culture vessel assemblies were connected to fluidic lines after sterilization and slotted into an eVOLVER Smart Sleeve for monitoring of OD and control of temperature and stir rate.

### PID algorithm development and tuning

To control the rate of dilution, we used the following equation to determine the percentage of selection media to add:

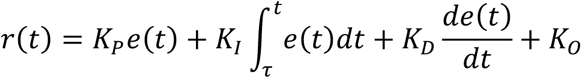

where *K*_*P*_, *K*_*I*_, *K*_*D*_, and *K*_*O*_, are empirically determined constant multipliers of proportional, integral, derivative, and offset terms, and *e*(*t*) is the difference between the actual growth rate and the target growth rate. To estimate *K*_*P*_, *K*_*I*_, *K*_*D*_, and *K*_*O*_, we used the the Ziegler-Nichols method^19^ for initially tuning the parameters with the pre-evolution strain, ZZ-Y323. *K*_*I*_ and *K*_*D*_ were first set to zero and *K*_*P*_ was increased until regular oscillations in growth rate were observed (Supplementary Fig. 2). This resulted in a *K*_*P*_ = 4.

Using the parameters obtained during the oscillation and the Ziegler-Nichols estimation:

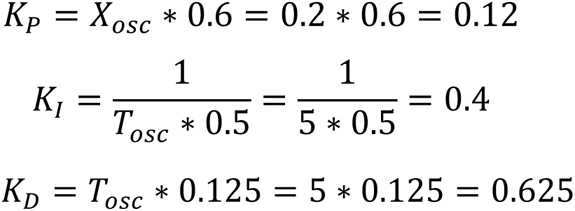

These initial values were empirically tuned to achieve the final values of *K*_*P*_ = 0.07, *K*_*I*_ = 0.05, *K*_*D*_ = 0.2 and *K*_*O*_ = 0.

These constants were then used to calculate *r*(*t*) at any given point during evolution. *r*(*t*) would then be used to determine the ratios of media to add during each dilution step by controlling the pump runtime. For example, if an *r*(*t*) = 0.25 was determined with a pump runtime of 5 seconds, the pump for the base media would run for [1 − *r*(*t*)] ∗ 5 seconds = 3.75 seconds while the pump for the full selection media would run for *r*(*t*) ∗ 5 seconds = 1.25 seconds.

The integral error 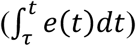 was reset at every instance the proportional error (*e*(*t*)) became negative, and the offset (*K*_*O*_) was updated to equal *r*(*t*) at that time. This was done to allow the PID controller to be more sensitive to the integral error and to avoid the bias that would result from the initial conditions having minimal selection pressure.

### *Pf*DHFR evolution

eVOLVER was set to morbidostat mode with the PID settings described above, a target doubling time of 8 hours, and one dilution step per hour. A culture of ZZ-Y435 was grown to saturation in SC-HW and then inoculated 1:50 in eVOLVER vials. SC-HW served as the base media, while SC-HW + 3 mM pyrimethamine served as the full selection media. (3mM was previously determined as the maximum soluble concentration of pyrimethamine in media^2^.) After inoculation, the eVOLVER PID script was initiated and evolution commenced. During evolution, the only user intervention was media exchange and periodic sampling of cultures. After 725 hrs, all cultures achieved growth rates near wild-type levels in the full selection condition (Supplementary Fig. 3), so the experiment was stopped and cultures were frozen in glycerol stocks.

### *Tm*HisA evolution

eVOLVER was set to morbidostat mode with the PID settings described above, a target doubling time of 8 hours, and one dilution step per hour. A culture of ZZ-Y323 was grown to saturation in SC-UL and then inoculated 1:50 in eVOLVER vials. SC-ULH + 7.76 mg/L (50 µM) histidine served as the base media, while SC-ULH served as the full selection media. After inoculation, the eVOLVER PID script was initiated and evolution commenced. During evolution, the only user intervention was media exchange and periodic sampling of cultures. After 715 hrs, all cultures achieved growth rates near wild-type levels in the full selection condition (Supplementary Fig. 4), so the experiment was stopped and cultures were frozen in glycerol stocks.

### Bulk DNA sequencing and characterization

Final evolution timepoints of *Pf*DHFR and *Tm*HisA were regrown in SC-HW and SC-ULH media, respectively, from glycerol stocks. The orthogonal plasmids encoding evolved *Pf*DHFR or *Tm*HisA were extracted from the bulk cultures as described above, PCR amplified, and sequenced via Sanger sequencing. Mutation frequencies were calculated from Sanger sequencing files with QSVanalyzer as previously described^2^. However, V1 from *Tm*HisA evolution could not be revived from the glycerol stock due to a stocking mistake and was not included for bulk DNA sequencing.

### *Tm*HisA isolated mutant cloning

Final evolution time-points of *Tm*HisA were streaked onto SC-ULH solid media. Individual colonies were regrown in SC-ULH media and the orthogonal plasmid DNA was extracted from the cultures as described above. The evolved *Tm*HisA sequences were sequenced and cloned into a nuclear *CEN6/ARS4* expression vector under control of the pRPL18B promoter and with the *LEU2* selection marker. Since each colony can have different *Tm*HisA mutants due to the multicopy nature of the orthogonal plasmid in OrthoRep, the cloned plasmids were sequenced again to determine the exact mutant of *Tm*HisA being characterized. The resulting plasmids were transformed into ZZ-Y354, which lacks *his6*, for growth rate measurements.

### *Tm*HisA growth rate measurements

Yeast strains containing each *Tm*HisA mutant, WT *Tm*HisA, *S. cerevisiae* HIS6, or none of the above expressed from a nuclear plasmid were grown to saturation in SC-L and diluted 1:100 in SC-LH. Three 100 uL replicates of each strain were placed into a 96 well clear-bottom tray, sealed, and grown at 30 °C. Cultures were continuously shaken and OD_600_ was measured every 30 minutes automatically for 15 hours (Tecan Infinite M200 Pro) according to a previously described protocol^20^. A custom MATLAB script (growthassayV3.m), included in supplemental files, was used to calculate growth rates from raw OD_600_ data. The script carries out a logarithmic transformation of the OD_600_ data. The linear region of the transformed data as a function of time corresponds to log phase growth. A sliding window approach is used to find and fit this linear region in order to calculate the doubling time during log phase growth. This doubling time (*T*) is converted to the continuous growth rate plotted in Fig. 3c by the formula *ln(2)*/*T*.

### Statistical analysis

Statistical analysis was done using GraphPad Prism and one-way ANOVA with multiple comparisons versus wild-type *Tm*HisA and corrected for multiple comparisons. Results are reported at p < 0.05.

## Supplementary Information

**Supplementary Figure 1.**
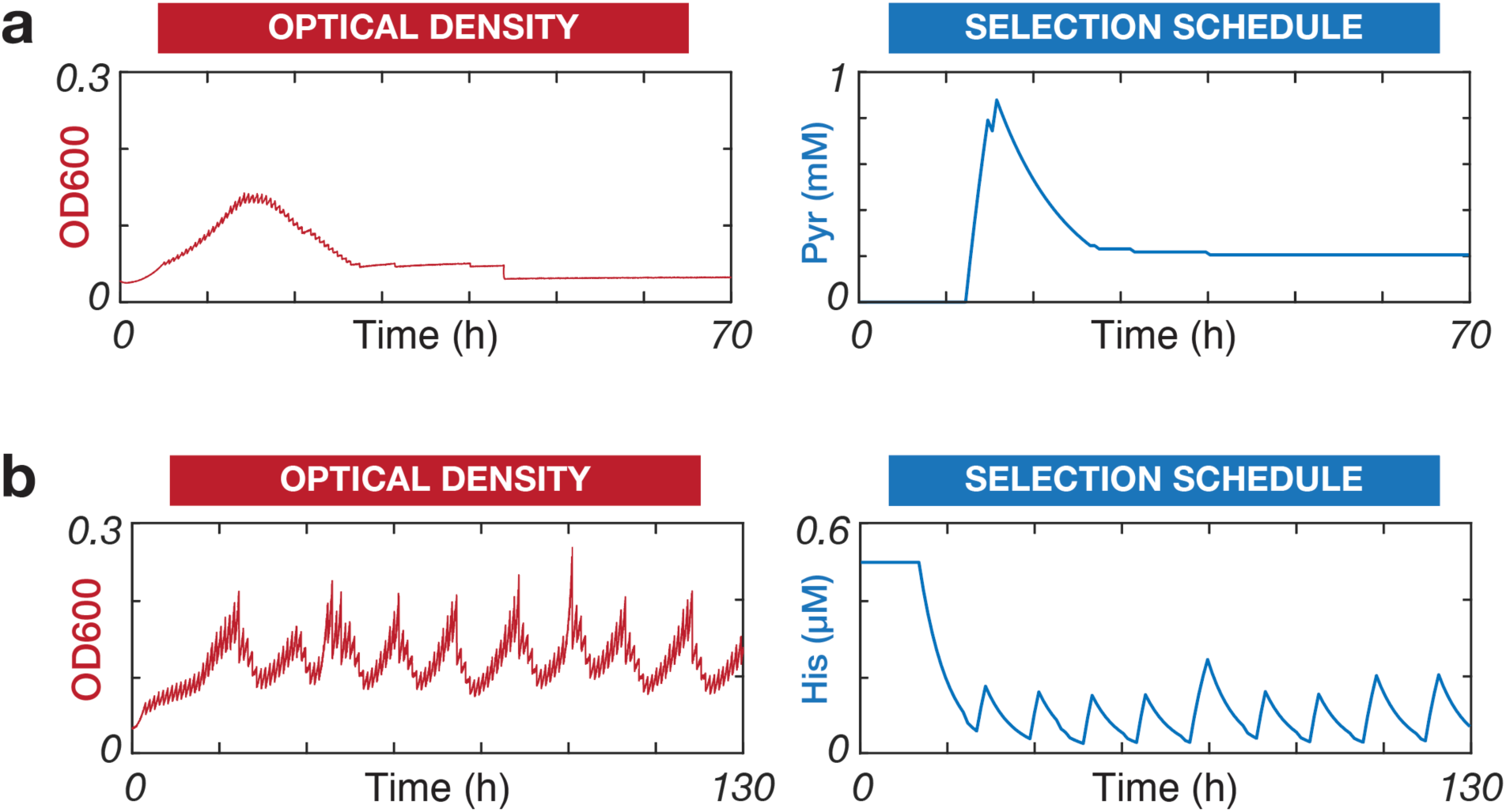
Sample evolution experiments in eVOLVER using the control algorithm derived from Toprak et al. 2013^15^. Briefly, this algorithm sampled OD at fixed intervals and if (1) the current OD is greater than a threshold and (2) if the current OD is greater than the previous OD, the growing cultures were diluted with the selection media. Otherwise, the culture would be diluted with the base media **(a**) An example of *Pf*DHFR evolution using the Toprak et al. algorithm. ZZ-Y435 was inoculated into eVOLVER and grown as described in **Methods** except the control algorithm was as described in Toprak et al.^15^. The OD of the adapting culture (red) and the calculated concentration of pyrimethamine (blue) are shown. After the initial increase and subsequent decrease of pyrimethamine, no further growth is seen in 40 hours. **(b**) An example of *Tm*HisA evolution using the Toprak et al.^15^ algorithm. ZZ-Y323 was inoculated into eVOLVER and grown as described in **Methods** except for the control algorithm. The OD of the adapting culture (red) and the calculated concentration of histidine in the media (blue) are shown. The concentration of histidine is observed to oscillate during selection and is unable to successfully adapt.

**Supplementary Figure 2.**
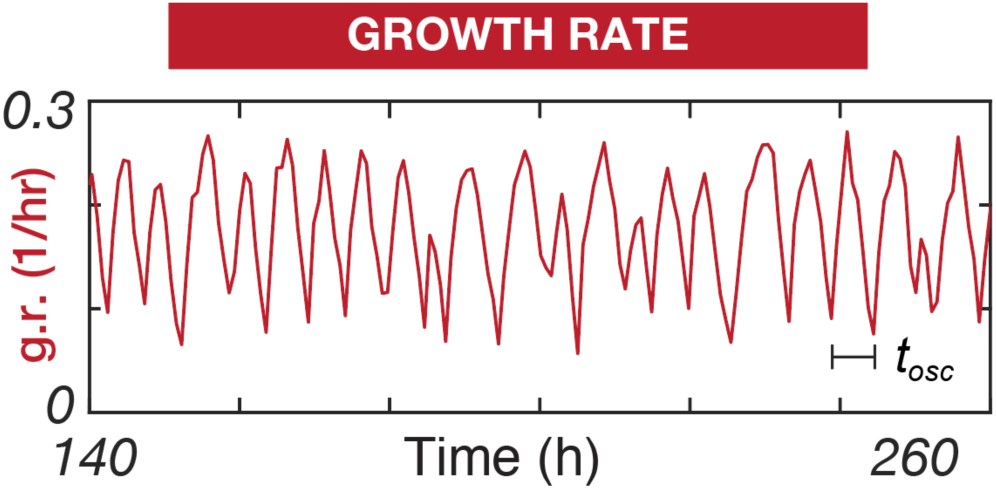
Oscillation of growth rate of ZZ-Y323 during the empirical determination of PID settings. ZZ-Y323 was inoculated into eVOLVER and SC-UL was used as the base media and SC-ULH was used as the full adaptation media. *K*_*P*_ was iterated from 0 to 4 over 120 hours while *K*_*I*_ and *K*_*D*_ were set to zero. Oscillations were observed when *K*_*P*_ = 4, and the period and amplitude of the oscillation were used to estimate the parameters for the PID control algorithm.

**Supplementary Figure 3.**
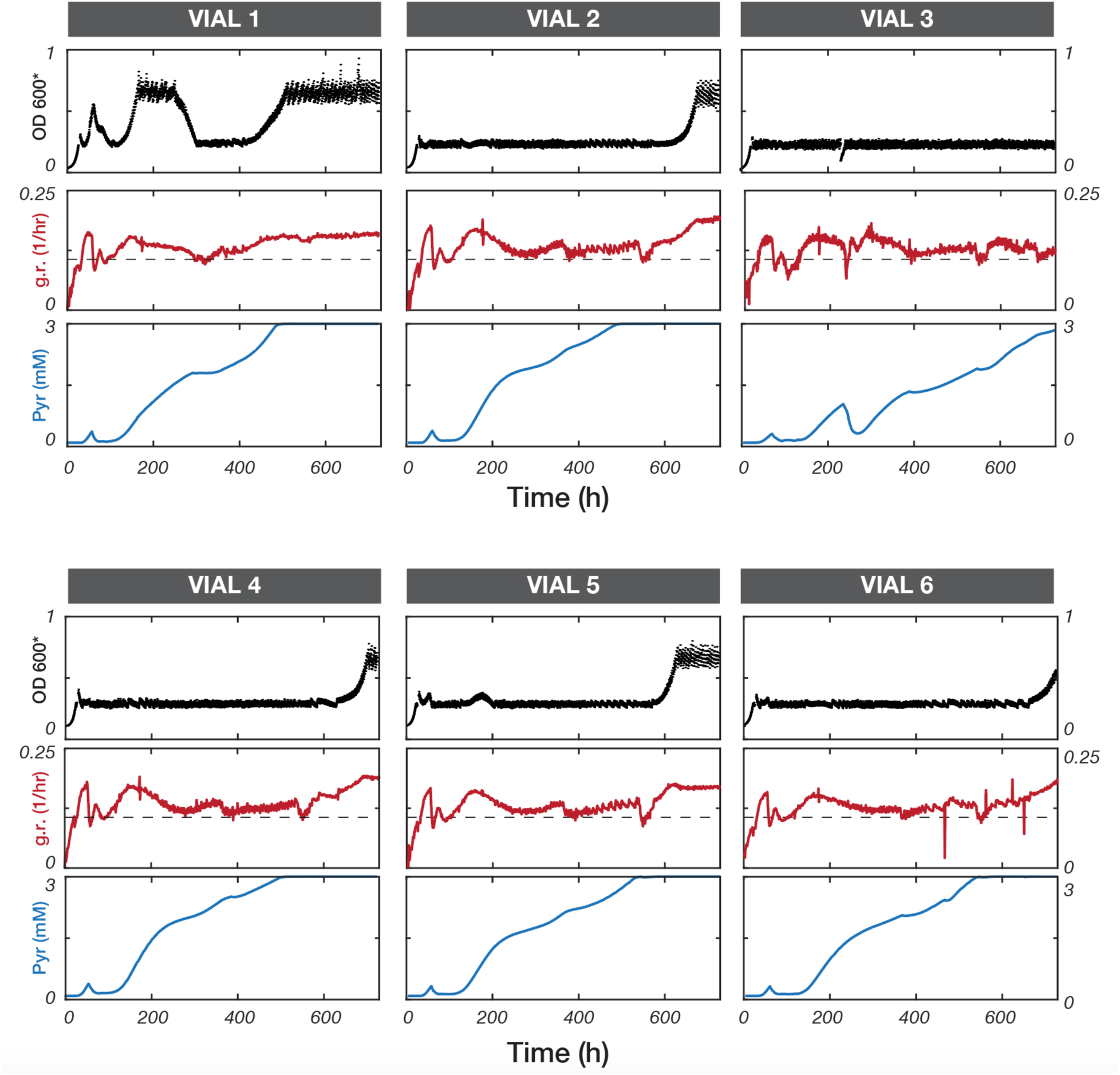
Adaptation history for all six replicates during *Pf*DHFR evolution. OD (black dot), growth rate (red line), target growth rate (black dash), and pyrimethamine concentration (blue line) are plotted for all six independent replicates. All six independent cultures were able to adapt to 3 mM pyrimethamine without the need for pre-programmed selection schedules or user intervention except to replenish media stocks.

**Supplementary Figure 4.**
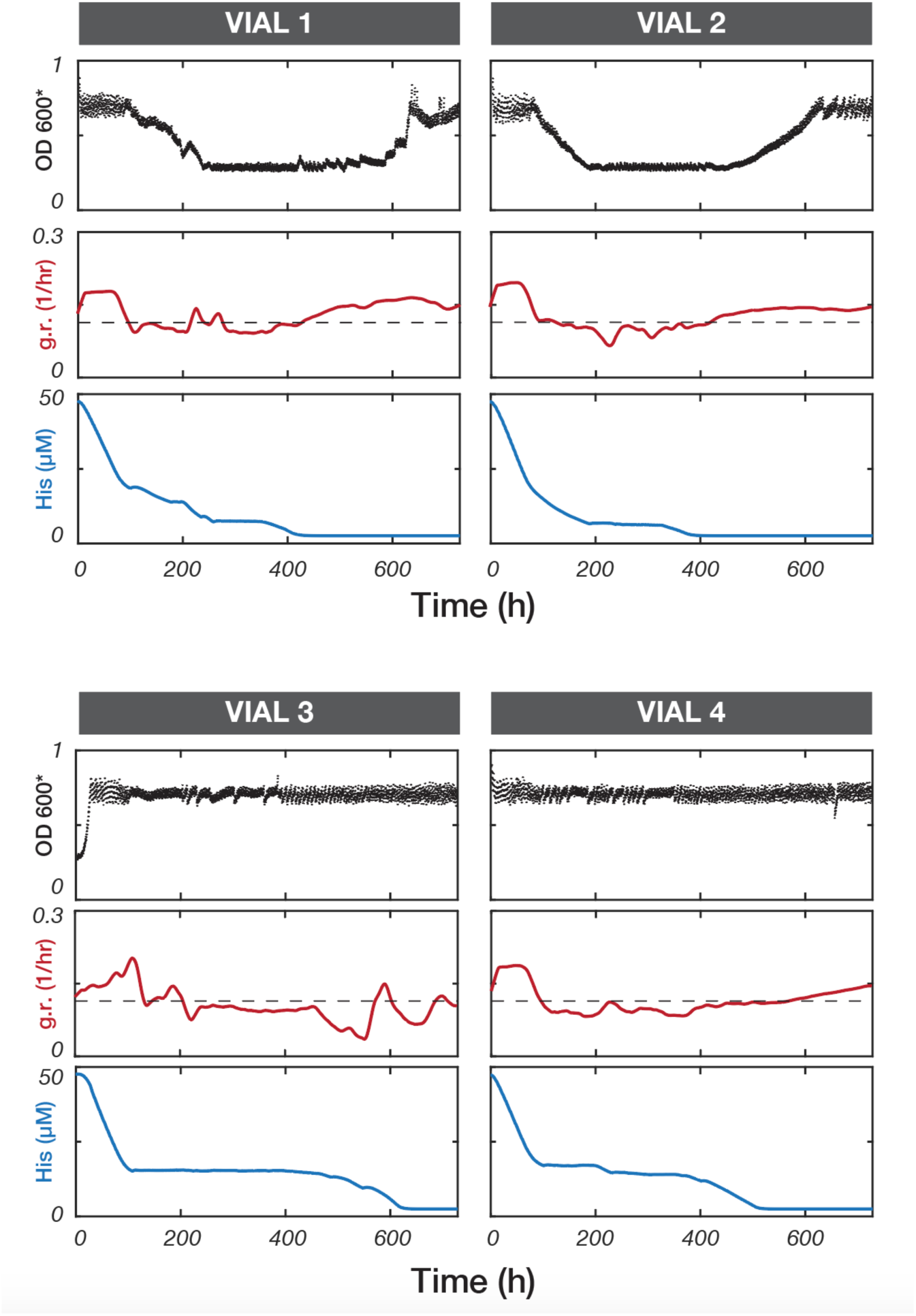
Adaptation history for all four replicates during *Tm*HisA evolution. OD (black dot), growth rate (red line), and histidine concentration (blue line) are plotted for all six independent replicates. All four cultures were able to adapt to media lacking histidine without the need for evolution schedules or user intervention except to provide eVOLVER with fresh base and full adaptation media in 700 hours of growth.

**Supplementary Table 1.**
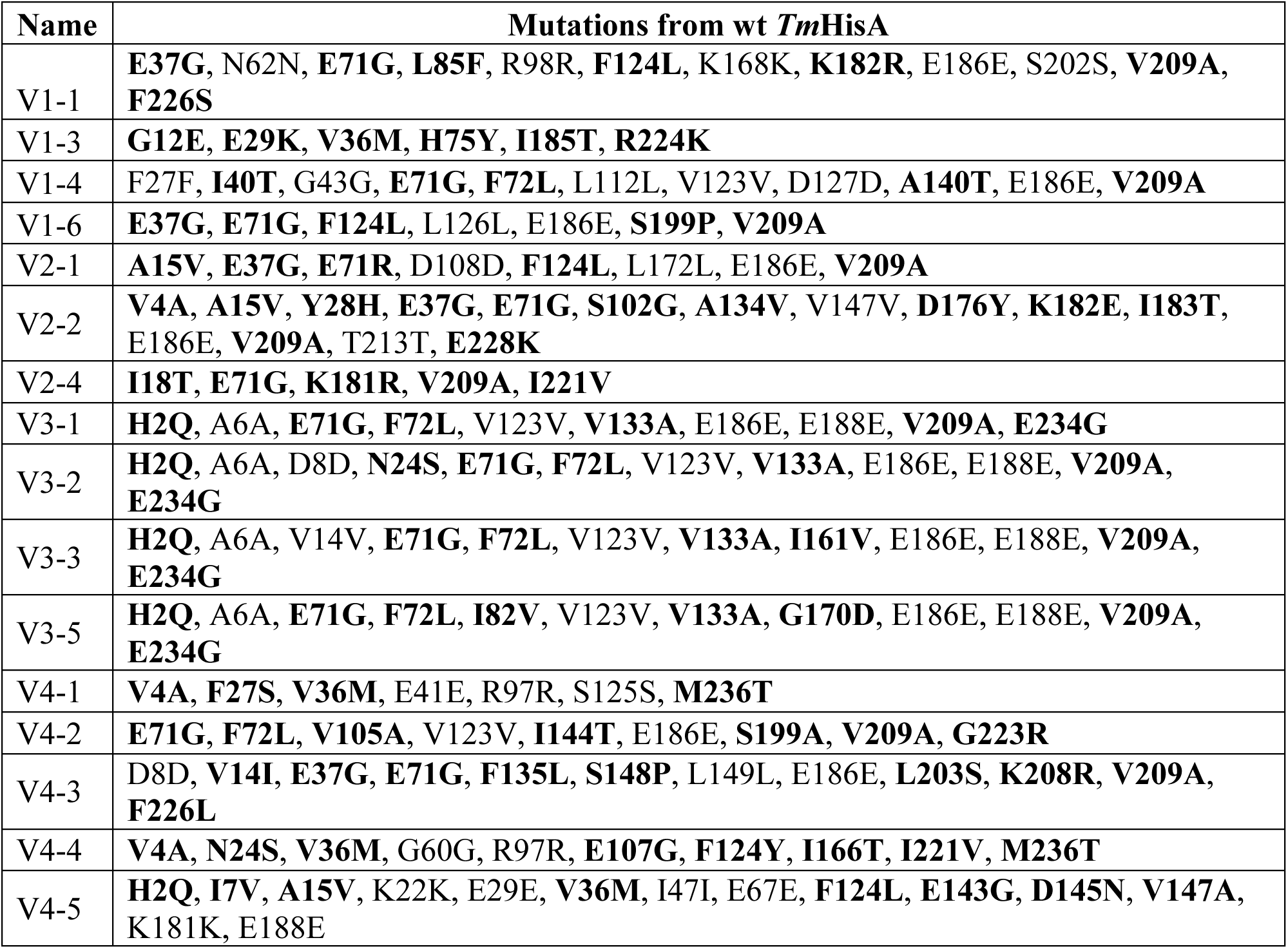
*Tm*HisA mutants characterized. Both nonsynonymous mutations (**bold**) and synonymous are noted. The order of these variants (top to bottom) corresponds to the order of the variants (left to right) in Figure 3c.

**Supplementary Table 2.**
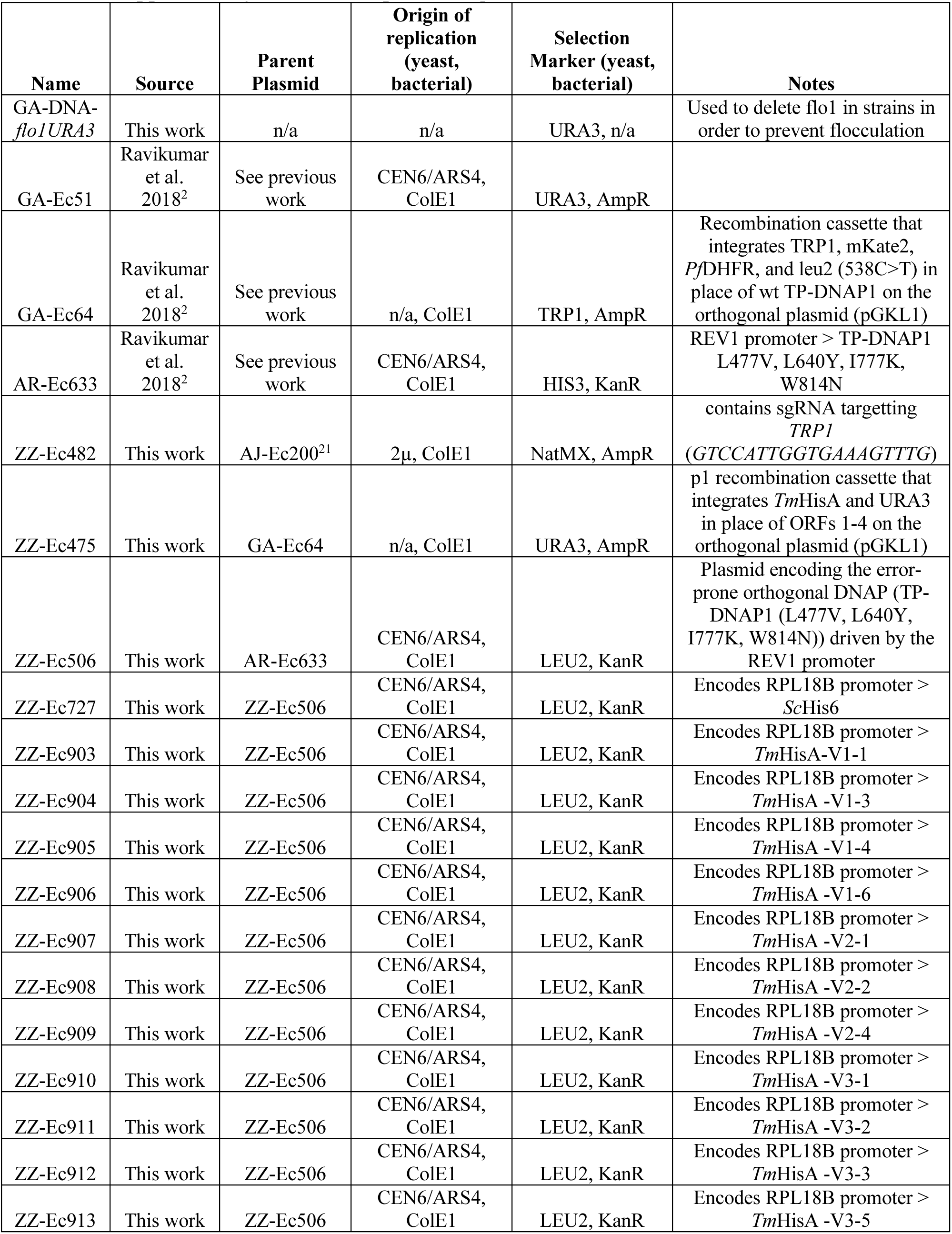
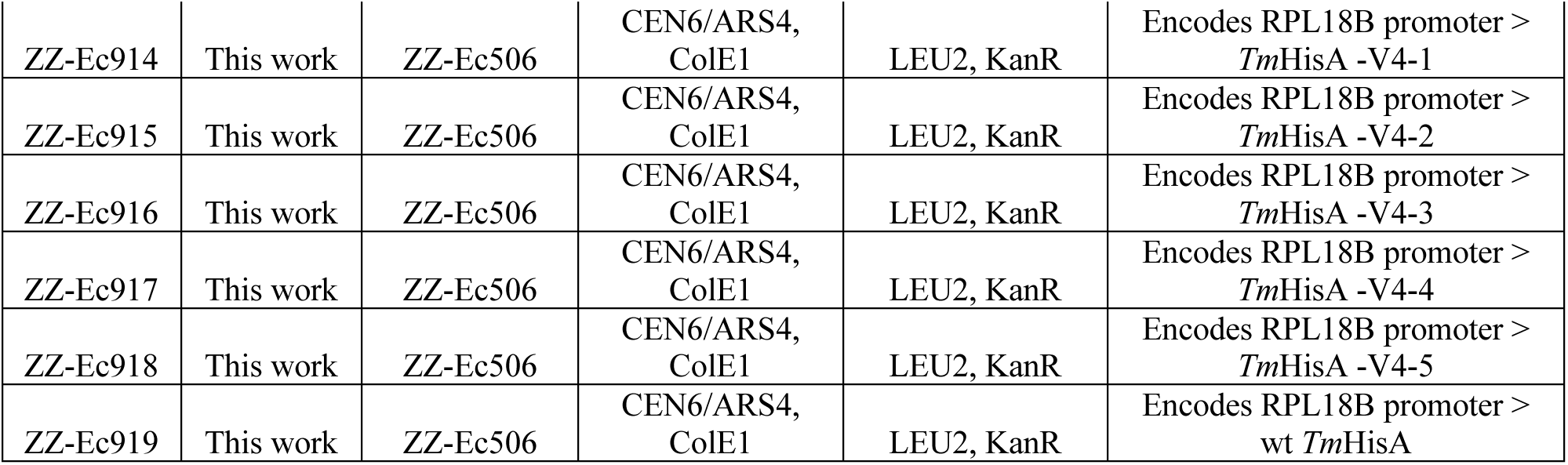
List of plasmids used in this study. For plasmids ZZ-Ec903 to ZZ-Ec918, see Supplementary Table 1 for specific sequences.

**Supplementary Table 3.**
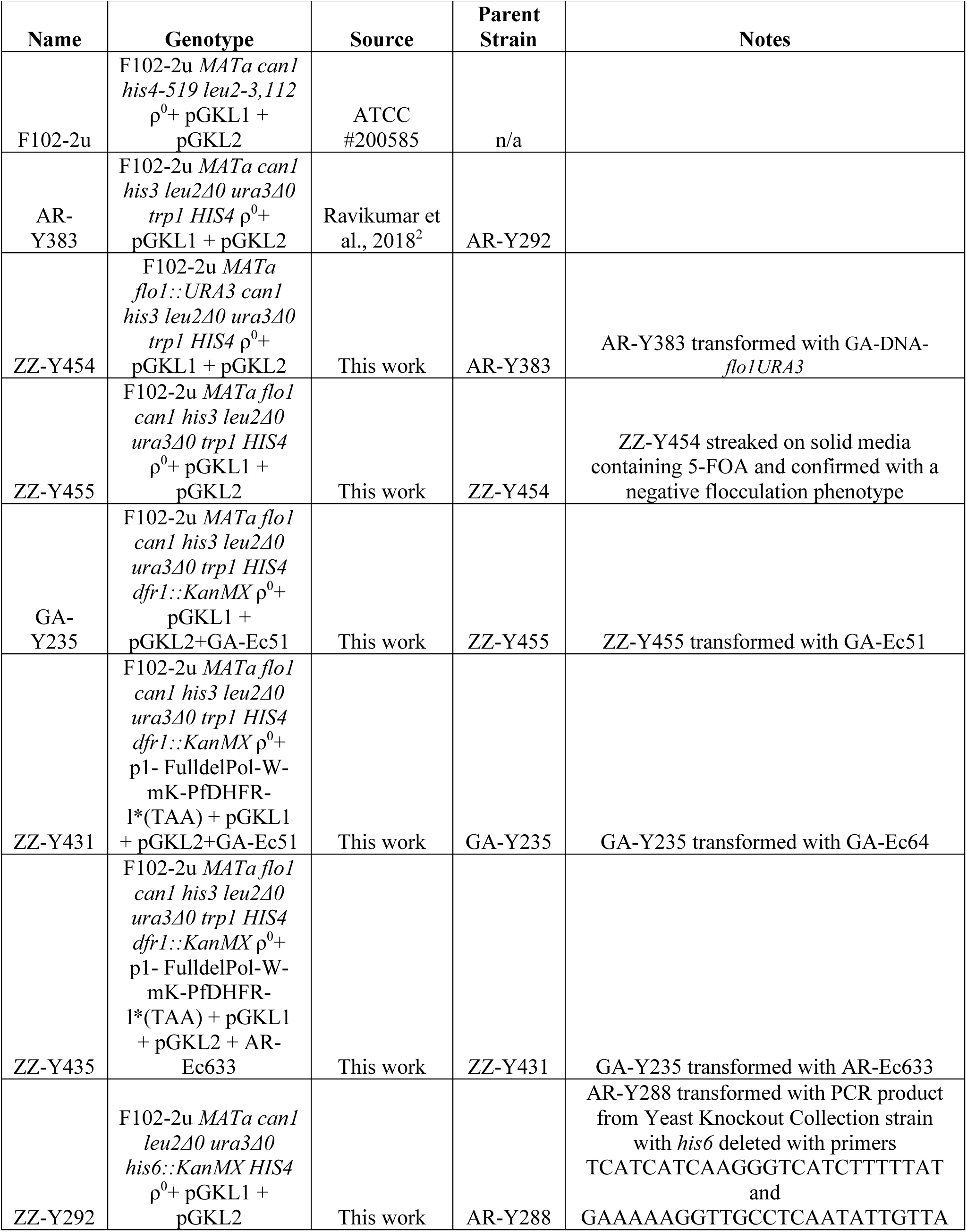
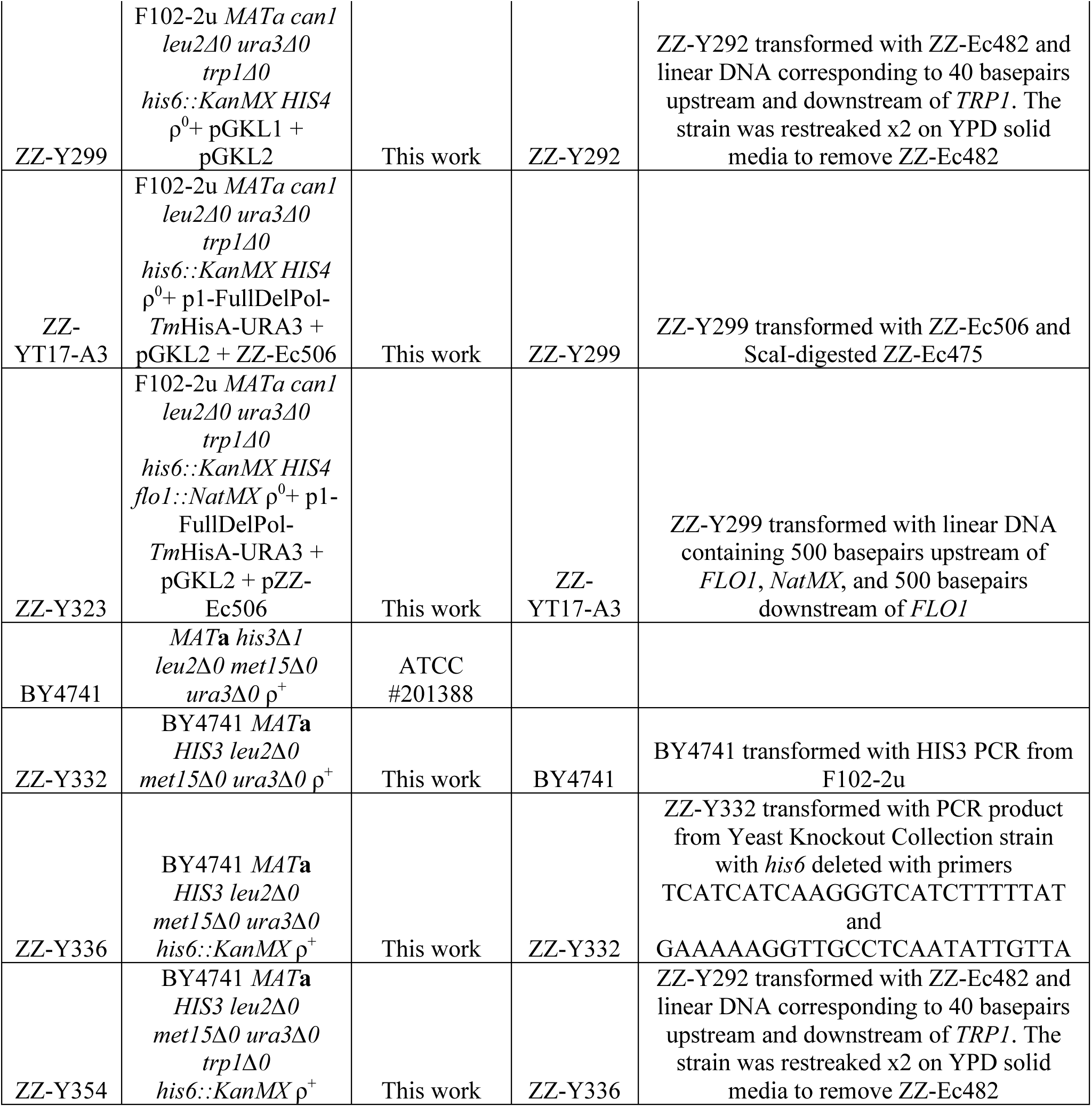
List of yeast strains used in this study.

