## Supplementary material for "Automated continuous evolution of proteins *in vivo*": Code associated with main paper: custom_script.py

```
import eVOLVER_module
import numpy as np
import os.path
import math
from scipy.optimize import curve_fit

###### Description of Code ##########
# This code is designed for eVOLVER to dynamically tune selection by altering the stringency of selection (AKA drug concentration)
# via diluting with a mixture of two choices of media; one media that is maximum drug concentration and one media that has no drug
# or a drug concentration at which selection first begins.
# Every 1 hour (or user definable timeline), a dilution event occurs to dilute a fixed amount based on the culture volume (an estimate),
# and the target growth rate (user defined). 

# PUMP TABLE
# PUMPS   VIAL    ROLE
# 0   0   NO DRUG
# 1   1   NO DRUG
# 2   2   NO DRUG
# 3   3   NO DRUG
# 4   4   NO DRUG
# 5   5   NO DRUG
# 6   6   NO DRUG
# 7   7   NO DRUG
# 8   0   DRUG
# 9   1   DRUG
# 10  2   DRUG
# 11  3   DRUG
# 12  4   DRUG
# 13  5   DRUG
# 14  6   DRUG
# 15  7   DRUG
# 16  0   WASTE
# 17  1   WASTE
# 18  2   WASTE
# 19  3   WASTE
# 20  4   WASTE
# 21  5   WASTE
# 22  6   WASTE
# 23  7   WASTE
# 24  12    NO DRUG
# 25  13    NO DRUG
# 26  12    DRUG
# 27  13    DRUG
# 28  12    WASTE
# 29  13    WASTE
# 30  N/A N/A
# 31  N/A N/A


def test (ODData, tempData, vials, elapsedTime, expName):
    # Set Stir rates
    MESSAGE = "13,10,15,10   ,10,8,7,10,  6,10,10,8,  10,5,10,10,"
    eVOLVER_module.stir_rate(MESSAGE)
        
    control = np.power(2,range(0,32))
    flow_rate = 1.0 #ml/sec

    # Global paramemters for PID
    VOLUME =  30.0                              # Volume assumed for dilution events (mL)
    DILUTION_OD = 0.25                          # OD at which dilutions start occurring
    MAX_OD = 0.75                               # High OD safeguard to prevent culture from leaving linear range
    DOUBLING_TIME = 8.0                         # User defined target doubling time
    TARGET_GR = math.log(2.0)/DOUBLING_TIME     # Calculated based off the user defined doubling time
    STEPS_PER_HOUR = 1                          # User defined number of dilution events per hour. Volumes are adjusted accordingly
    DILUTION_RATE = float(VOLUME / DOUBLING_TIME) # mL / hour
    VOLUME_PER_DILUTION = float(DILUTION_RATE/ STEPS_PER_HOUR)
    TIME_BETWEEN_DILUTIONS = float(60/STEPS_PER_HOUR) #minutes

    # PID Parameters
    # Kp, Ki, Kd may need to be changed for different selections
    Kp =         [0.07, 0.07,        0.07, 0.07,         0.07, 0.07,          0.07, 0.07] 
    Ki =         [0.05, 0.05,        0.05, 0.05,     0.05, 0.05,      0.05, 0.05] 
    Kd =         [0.2, 0.2,        0.2, 0.2,         0.2, 0.2,          0.2, 0.2] 
    PIDOffset = [0.0, 0.0,    0.0, 0.0,     0.0, 0.0,      0.0, 0.0]

    if elapsedTime > 1: # initial time for growth, hours

        #Load pump log (all recorded in vial 0 log across all 16, since pump time occurs simultaneously for all)
        savePath = os.path.dirname(os.path.realpath(__file__))
        filePath =  "%s/%s/pump_log/vial0_pump_log.txt" % (savePath,expName)
        data = np.genfromtxt(filePath, delimiter=',')
        lastPump = data[len(data)-1][0]

        # if enough time passed for the next dilution to occur
        if ((elapsedTime- lastPump)*3600) > (TIME_BETWEEN_DILUTIONS * 60): # convert to seconds and compare

            # determine which pumps to fire (e.g. drug or no drug) based on current OD
            cleanCommand = 0;

            for x in range(0,4):
                pumpCommand = 0
                cleanCommand = cleanCommand + control[x+16]

                PIDPumpCommandNoDrug = 0
                PIDPumpCommandDrug   = 0

                # 1. Reads GR from last dilution time
                GRPath =  "%s/%s/growth_rate/vial%d_growth_rate.txt" % (savePath,expName,x)
                growthRateData = np.genfromtxt(GRPath, delimiter=',')
                lastGR = growthRateData[len(growthRateData)-1][1]

                # 2. Reads drug concentration from last dilution time
                drugLogPath =  "%s/%s/drug_log/vial%d_drug_log.txt" % (savePath,expName,x)
                drugLogData = np.genfromtxt(drugLogPath, delimiter=',')
                drugConc = drugLogData[len(drugLogData)-1][1]

                # 3. Read and calculates the current OD (average over 5 measurements)
                ODPath =  "%s/%s/OD/vial%d_OD.txt" % (savePath,expName,x)
                ODData = np.genfromtxt(ODPath, delimiter=',')
                
                # 4. Reads PID parameters from last round
                PIDPath = "%s/%s/PIDLog/vial%d_PIDLog.txt" % (savePath,expName,x)
                PIDData = np.genfromtxt(PIDPath, delimiter=',')
                lastIError = PIDData[len(PIDData)-1][2]
                lastPError = PIDData[len(PIDData)-1][1]
                lastVialPump = PIDData[len(PIDData)-1][0]

                # 5. Reads the offset parameter from last round
                offsetLogPath = "%s/%s/offset_log/vial%d_offset.txt" % (savePath,expName,x)
                offsetLogData = np.genfromtxt(offsetLogPath, delimiter=',')
                if len(offsetLogData) > 2:
                    PIDOffset[x] = offsetLogData[len(offsetLogData)-1][1]

                # CALCULATIONS #
                # Calculates OD for a vial (across 5 measurements)
                averageOD = 0
                for n in range(1,6):
                    averageOD = averageOD + (ODData[len(ODData)-n][1]/5)

                # Calculates OD after previous dilution (across 5 measurements)
                ODAfterDilution = 0
                for n in range(1,6):
                    ODAfterDilution = ODAfterDilution + (ODData[len(ODData)-n-int((360/STEPS_PER_HOUR))+18][1]/5)

                # Cleans up OD readings for growth rate calculation
                # This only occurs of OD < 0
                if ODAfterDilution < 0 or averageOD < 0:
                    growthRate = 0
                    growthRate_OLD = 0
                    ODFitGuess = 0
                else:
                    # Calculates growth rate
                    ODWindow =   ODData[len(ODData)-int((360/STEPS_PER_HOUR))+15:,1]
                    timeWindow = ODData[len(ODData)-int((360/STEPS_PER_HOUR))+15:,0]
                    timeWindow = timeWindow - timeWindow[0]

                    # Fits the OD and time to a exponential curve, func
                    popt, pcov = curve_fit(func,timeWindow,ODWindow,bounds=([ODAfterDilution - 0.05, 0],[ODAfterDilution + 0.05, 0.5]))
                    
                    growthRate = popt[1]

                # If the vial has been below the dilution OD for 4 rounds, and the culture has been growning for a while:
                # Dilute with no drug media to wash out drug. 
                # Otherwise, dilute normally 
                if (elapsedTime - lastVialPump) > (TIME_BETWEEN_DILUTIONS*4/60) and drugConc > 0.05 and elapsedTime > 144:
                    print ("!!!!RESCUE DILUTION SHOULD HAVE OCCURRED!!!!")
                    pumpCommand = pumpCommand + control[x] + control[x+16]
                    MESSAGE   = "%s,0,%f," % ("{0:b}".format(pumpCommand) , 3)
                    eVOLVER_module.fluid_command(MESSAGE, x, elapsedTime, TIME_BETWEEN_DILUTIONS *60, expName, 3, 'n')
                    pidFile = open(PIDPath, "a+")
                    pidFile.write("%f,%f,%f,%f,%f,%f,%f,%f,%f\n" % (elapsedTime, 0, 0, 0, 0, 0, 0, 0, 0))
                    pidFile.close()

                    drugConc = (VOLUME * drugConc)/(VOLUME + 3)
                    drugFile = open(drugLogPath,"a+")
                    drugFile.write("%f,%f\n" %  (elapsedTime, drugConc))
                    drugFile.close()
                elif averageOD > DILUTION_OD:
                    # PID Calculations
                    pError = 0
                    iError = 0
                    dError = 0
                    MESSAGEDrug = "0,0,0"
                    MESSAGENoDrug = "0,0,0"
                    PIDControl = 0

                    # Calculates proportional error, e(t)
                    pError = growthRate - TARGET_GR

                    # Calculates Integrate[e(t), dt, t0, t] with trapezoid rule
                    # The integral is calculated every time the proportional 
                    # error switches signs (so that every time that e[t] = 0, the integral term resets)
                    if pError == 0 or lastPError / pError < 0:
                        iError = (growthRate + lastGR  - 2*TARGET_GR) / STEPS_PER_HOUR
                    else:
                        iError = lastIError + (growthRate + lastGR  - 2*TARGET_GR) / STEPS_PER_HOUR

                    # Calculates d e(t)/dt
                    dError = (growthRate - lastGR) * STEPS_PER_HOUR

                    # If the machine has been acting normally, if the growth rate dips below the target growth rate,
                    # a new offset is calculated and used for future rounds of control
                    if elapsedTime > 72 and pError < 0 and lastPError > 0:
                        print ("OFFSET LOG DIAGNOSTIC elapsedTime: %f" %(elapsedTime))
                        print ("OFFSET LOG DIAGNOSTIC pError: %f" %(pError))
                        print ("OFFSET LOG DIAGNOSTIC lastPError: %f" %(lastPError))
                        print ("OFFSET LOG DIAGNOSTIC len(drugLogData): %f" %(len(drugLogData)))
                        newPIDoffset = 0
                        for n in range(6,11):
                            newPIDoffset = newPIDoffset + (drugLogData[len(drugLogData)-n][1])/5
                        PIDOffset[x] = newPIDoffset
                        offset_file = open(offsetLogPath, "a+")
                        offset_file.write("%f, %f\n" % (elapsedTime, newPIDoffset))
                        offset_file.close()

                    # Calculates the drug dilution percentage
                    PIDControl = PIDOffset[x] + pError * Kp[x] + iError * Ki[x] + dError * Kd[x]
                    
                    # PID Pump Command
                    PIDPumpCommandNoDrug = PIDPumpCommandNoDrug + control[x] + control[x+16]
                    PIDPumpCommandDrug   = PIDPumpCommandDrug   + control[x+8] + control[x+16]

                    # Cleans up the controls in case they were calculated to be greater than 1 or less than 0
                    if PIDControl > 1:
                        PIDControl = 1
                        PIDPumpCommandNoDrug = 0
                    elif PIDControl < 0:
                        PIDControl = 0
                        PIDPumpCommandDrug = 0

                    # Option to dilute twice if the OD is > MAX_OD. This is to keep the culture within the linear range of the detectors
                    numDils = 1
                    if averageOD > MAX_OD:
                        numDils = 2
                        print ("Double Dilution Occurred; numDils = 2")

                    # Sends dilution command to RaspberryPi
                    MESSAGENoDrug = "%s,0,%f," % ("{0:b}".format(PIDPumpCommandNoDrug) , numDils * VOLUME_PER_DILUTION/flow_rate * ( 1 - PIDControl))
                    MESSAGEDrug   = "%s,0,%f," % ("{0:b}".format(PIDPumpCommandDrug) , numDils * VOLUME_PER_DILUTION/flow_rate * PIDControl)
                    if PIDControl < 1:
                        eVOLVER_module.fluid_command(MESSAGENoDrug, x, elapsedTime, TIME_BETWEEN_DILUTIONS *60, expName, numDils * VOLUME_PER_DILUTION/flow_rate * ( 1 - PIDControl), 'n')
                        print ("Message NoDrug: %s" %(MESSAGENoDrug))
                    if PIDControl > 0:
                        eVOLVER_module.fluid_command(MESSAGEDrug, x, elapsedTime, TIME_BETWEEN_DILUTIONS *60, expName, numDils * VOLUME_PER_DILUTION/flow_rate * PIDControl, 'n')
                        print ("Message Drug: %s" %(MESSAGEDrug))

                    # Calculates the new drug concentration of the vial
                    drugConc = (VOLUME * drugConc + PIDControl * numDils * VOLUME_PER_DILUTION/flow_rate)/(VOLUME + numDils * VOLUME_PER_DILUTION/flow_rate)

                    # Updates the PID parameter file
                    pidFile = open(PIDPath, "a+")
                    pidFile.write("%f,%f,%f,%f,%f,%f,%f,%f,%f\n" % (elapsedTime, pError, iError, dError, Kp[x] * pError, Ki[x] * iError, Kd[x] * dError,PIDOffset[x], PIDControl))
                    pidFile.close()


                # 1. Update GR file with current GR
                GRFile = open(GRPath,"a+")
                GRFile.write("%f,%s\n" %  (elapsedTime, growthRate))
                GRFile.close()

                # 2. Updates drug concentration file only if a dilution event happened
                if averageOD > DILUTION_OD:
                    drugFile = open(drugLogPath,"a+")
                    drugFile.write("%f,%f\n" %  (elapsedTime, drugConc))
                    drugFile.close()

                # 3. Updates a log file with an assortment of values for plotting
                logPath = "%s/%s/logs/vial%d_log.txt" % (savePath,expName,x)
                logFile = open(logPath, "a+")
                logFile.write ("Vial %d\n" %(x))
                logFile.write ("Average OD: %f\n" %(averageOD))
                logFile.write ("Target Growth Rate: %f\n" %(TARGET_GR))
                logFile.write ("Growth Rate: %f\n" %(growthRate))
                logFile.write ("pError: %f\n" %(pError))
                logFile.write ("i error: %f\n" %(iError))
                logFile.write ("dError: %f\n" %(dError))
                logFile.write ("PID Offset: %f\n" %(PIDOffset[x]))
                logFile.write ("PIDControl: %f\n" %(PIDControl))
                logFile.write ("Message Drug: %s\n" %(MESSAGEDrug))
                logFile.write ("Message NoDrug: %s\n\n" %(MESSAGENoDrug))
                logFile.close()


                # Terminal display values for each vial:
                print ("Vial %d" %(x))
                print ("Average OD: %f" %(averageOD))
                print ("Target Growth Rate: %f" %(TARGET_GR))
                print ("Growth Rate: %f" %(growthRate))
                print ("last p error: %f" %(lastPError))
                print ("pError: %f" %(pError))
                print ("i error: %f" %(iError))
                print ("dError: %f" %(dError))
                print ("PID Offset: %f" %(PIDOffset[x]))
                print ("PIDControl: %f" %(PIDControl))
                
                # Vial13_logFile accounts for all Terminal output
                logPath = "%s/%s/logs/vial13_log.txt" % (savePath,expName)
                logFile = open(logPath, "a+")
                logFile.write ("Elapsed Time: %f" %(elapsedTime))
                logFile.write ("Vial %d\n" %(x))
                logFile.write ("Average OD: %f\n" %(averageOD))
                logFile.write ("Target Growth Rate: %f\n" %(TARGET_GR))
                logFile.write ("Growth Rate: %f\n" %(growthRate))
                logFile.write ("pError: %f\n" %(pError))
                logFile.write ("i error: %f\n" %(iError))
                logFile.write ("dError: %f\n" %(dError))
                logFile.write ("PID Offset: %f\n" %(PIDOffset[x]))
                logFile.write ("PIDControl: %f\n" %(PIDControl))
                logFile.write ("Message Drug: %s\n" %(MESSAGEDrug))
                logFile.write ("Message NoDrug: %s\n\n" %(MESSAGENoDrug))
                logFile.close()

            # Runs output pumps for 5 seonds to ensure volume of vial is constant
            CLEAN_MESSAGE = "%s,0,5," % ("{0:b}".format(cleanCommand))
            print ("CLEAN_MESSAGE: %s" %(CLEAN_MESSAGE))
            eVOLVER_module.fluid_command(CLEAN_MESSAGE, 0, elapsedTime, TIME_BETWEEN_DILUTIONS *60, expName, 5,'y')


def func(x, a, b):
    # Exponential function for fit
    return a * np.exp(b * x)


def measure_growthRate(x, savePath, expName):
    ## Load files from proper directories
    ODPath =  "%s/%s/OD/vial%d_OD.txt" % (savePath,expName,x)
    pump_path =  "%s/%s/pump_log/vial0_pump_log.txt" % (savePath,expName)
    ODData = np.genfromtxt(ODPath, delimiter=',')
    pump_data = np.genfromtxt(pump_path, delimiter=',')

    ### Gets time frame from last diution event to current event
    time_buffer = 3 #minutes
    growth_start= pump_data[len(pump_data)-2][0] + (time_buffer/60) # hours
    growth_end = pump_data[len(pump_data)-1][0]

    ## Gets part of data that is part of the growth curve
    segmented_OD = ODData[np.where(np.logical_and(ODData >=growth_start, ODData<=growth_end))[0]]

    ## Averages the OD data from start of curve and end of curve
    averaged_values = 10
    OD_start = np.mean(segmented_OD[0:averaged_values,1])
    OD_end = np.mean(segmented_OD[len(segmented_OD)-averaged_values :len(segmented_OD),1])

    ## Calculates growth rate
    growth_rate = ((OD_end-OD_start)/OD_start)/(growth_end - growth_start )

    return growth_rate
```
