## Supplementary material for "Automated continuous evolution of proteins *in vivo*": Code associated with main paper: eVOLVER_module.py

```
import socket
import os.path
import shutil
import time
import pickle
import numpy as np
import numpy.matlib
import matplotlib
import scipy.signal
matplotlib.use('Agg')
import matplotlib.pyplot as plt
plt.ioff()

def read_OD(vials):
    od_cal = np.genfromtxt("OD_cal.txt", delimiter=',')
    UDP_IP = "192.168.1.2"
    UDP_PORT = 5554
    MESSAGE='2125,2125,2125,2125,2125,2125,2125,2125,2125,2125,2125,2125,2125,2125,2125,2125,'
    sock = socket.socket(socket.AF_INET, # Internet
                     socket.SOCK_DGRAM) # UDP
    sock.settimeout(5)
    try:
        sock.sendto(MESSAGE, (UDP_IP, UDP_PORT))
        data, addr = sock.recvfrom(1024)
        data = data.split(',')
        for x in vials:
            try:
                data[x] = np.real(od_cal[2,x] - ((np.log10((od_cal[1,x]-od_cal[0,x])/(float(data[x]) - od_cal[0,x])-1))/od_cal[3,x]))
            except ValueError:
                print "OD Read Error"
                data = 'empty'
                break
    except socket.timeout:
        print "UDP Timeout (OD)"
        data = 'empty'

    return data

def update_temp(vials,exp_name):
    temp_cal = np.genfromtxt("temp_calibration.txt", delimiter=',')
    save_path = os.path.dirname(os.path.realpath(__file__))
    MESSAGE = ""
    for x in vials:
        file_path =  "%s/%s/temp_config/vial%d_tempconfig.txt" % (save_path,exp_name,x)
        data = np.genfromtxt(file_path, delimiter=',')
        temp_set = data[len(data)-1][1]
        temp_set = int((temp_set - temp_cal[1][x])/temp_cal[0][x])
        MESSAGE += str(temp_set)
        MESSAGE += ","
        
    UDP_IP = "192.168.1.2"
    UDP_PORT = 5553
    sock = socket.socket(socket.AF_INET, # Internet
                     socket.SOCK_DGRAM) # UDP
    sock.settimeout(5)
    sock.sendto(MESSAGE, (UDP_IP, UDP_PORT))
    try:
        data, addr = sock.recvfrom(1024)
        data = data.split(',')
        for x in vials:
            try:
                data[x] = (float(data[x]) * temp_cal[0][x]) + temp_cal[1][x]
            except ValueError:
                print "Temp Read Error"
                data = 'empty'
                break
    except socket.timeout:
        data = 'empty'
    return data

def fluid_command(MESSAGE, vial, elapsed_time, pump_wait, exp_name, time_on, file_write):
    UDP_IP = "192.168.1.2"
    UDP_PORT = 5552
    sock = socket.socket(socket.AF_INET, # Internet
                     socket.SOCK_DGRAM) # UDP
    sock.settimeout(5)
    save_path = os.path.dirname(os.path.realpath(__file__))
    file_path =  "%s/%s/pump_log/vial%d_pump_log.txt" % (save_path,exp_name,vial)
    data = np.genfromtxt(file_path, delimiter=',')
    last_pump = data[len(data)-1][0]
    if ((elapsed_time- last_pump)*3600) >pump_wait:
        sock.sendto(MESSAGE, (UDP_IP, UDP_PORT))
        if file_write == 'y':
            text_file = open(file_path,"a+")
            text_file.write("%f,%s\n" %  (elapsed_time, time_on))
            text_file.close()

def stir_rate (MESSAGE):
    UDP_IP = "192.168.1.2"
    UDP_PORT = 5551
    sock = socket.socket(socket.AF_INET, # Internet
                     socket.SOCK_DGRAM) # UDP
    sock.settimeout(5)
    sock.sendto(MESSAGE, (UDP_IP, UDP_PORT))

    

def parse_data(data, elapsed_time, vials, exp_name, file_name):
    save_path = os.path.dirname(os.path.realpath(__file__))
    if data == 'empty':
        print "%s Data Empty! Skipping data log..." % file_name
    else:
        for x in vials:
            file_path =  "%s/%s/%s/vial%d_%s.txt" % (save_path,exp_name,file_name,x,file_name)
            text_file = open(file_path,"a+")
            text_file.write("%f,%s\n" %  (elapsed_time, data[x]))
            text_file.close()

def initialize_exp(exp_name, vials):
    save_path = os.path.dirname(os.path.realpath(__file__))
    dir_path =  "%s/%s" % (save_path,exp_name)
    exp_continue = raw_input('Continue from exisiting experiment? (y/n): ')
    if exp_continue == 'n':
        OD_read = read_OD(vials)
        exp_blank = raw_input('Callibrate vials to blank? (y/n): ')
        pump_init = raw_input('Turn off pumps for this experiment? (y/n)')
        if exp_blank == 'y':
            OD_initial = OD_read
        else: 
            OD_initial = np.zeros(len(vials))
                    
        start_time = time.time()
        if os.path.exists(dir_path):
            shutil.rmtree(dir_path)
        os.makedirs("%s/OD" % dir_path)
        os.makedirs("%s/temp" % dir_path)
        os.makedirs("%s/temp_graph" % dir_path)
        os.makedirs("%s/pump_log" % dir_path)
        os.makedirs("%s/temp_config" % dir_path)
        os.makedirs("%s/growth_rate" % dir_path)
        os.makedirs("%s/ODset" % dir_path)

        os.makedirs("%s/drug_log" % dir_path)
        os.makedirs("%s/logs" % dir_path)
        os.makedirs("%s/PIDLog" % dir_path)
        os.makedirs("%s/offset_log" % dir_path)

        for x in vials:
            OD_path =  "%s/OD/vial%d_OD.txt" % (dir_path,x)
            text_file = open(OD_path,"w").close()
			
            growth_path =  "%s/growth_rate/vial%d_growth_rate.txt" % (dir_path,x)
            text_file = open(growth_path,"w")
            text_file.write("0,0\n0,0\n")
            text_file.close()
			
            temp_path =  "%s/temp/vial%d_temp.txt" % (dir_path,x)
            text_file = open(temp_path,"w").close()
			
            tempconfig_path =  "%s/temp_config/vial%d_tempconfig.txt" % (dir_path,x)
            text_file = open(tempconfig_path,"w")
            text_file.write("Experiment: %s vial %d, %r\n" % (exp_name, x, time.strftime("%c")))
            text_file.write("0,30\n")
            text_file.close()
			
            pump_path =  "%s/pump_log/vial%d_pump_log.txt" % (dir_path,x)
            text_file = open(pump_path,"w")
            text_file.write("Experiment: %s vial %d, %r\n" % (exp_name, x, time.strftime("%c")))
            if pump_init == 'y':
                text_file.write("99999999999999,0\n")
            else:
                text_file.write("0,0\n0,0\n")
            text_file.close()

            ODset_path =  "%s/ODset/vial%d_ODset.txt" % (dir_path,x)
            text_file = open(ODset_path,"w")
            text_file.write("Experiment: %s vial %d, %r\n" % (exp_name, x, time.strftime("%c")))
            text_file.write("0,0\n")
            text_file.close()
			
            drug_log_path =   "%s/drug_log/vial%d_drug_log.txt" % (dir_path,x)
            text_file = open(drug_log_path,"w")
            text_file.write("0,0\n0,0\n")
            text_file.close()
			
            log_path = "%s/logs/vial%d_log.txt" % (dir_path,x)
            text_file = open(log_path,"w")
            text_file.write("0,0\n0,0\n")
            text_file.close()

            log_path = "%s/PIDLog/vial%d_PIDLog.txt" % (dir_path,x)
            text_file = open(log_path,"w")
            #Log file consists of t, e(t), Int(e(t) dt, 0,t), d(e(t)/dt), Kp, Ki, Kd, output
            text_file.write("0,0,0,0,0,0,0,0,0\n")
            text_file.write("0,0,0,0,0,0,0,0,0\n")
            text_file.close()

            log_path = "%s/offset_log/vial%d_offset.txt" % (dir_path,x)
            text_file = open(log_path,"w")
            text_file.write("0,0\n0,0\n")
            text_file.close()
    else:
        pickle_path =  "%s/%s/%s.pickle" % (save_path,exp_name,exp_name)
        with open(pickle_path) as f:
            loaded_var  = pickle.load(f)
        x = loaded_var
        start_time = x[0]
        OD_initial = x[1]
    return start_time, OD_initial

def save_var(exp_name, start_time, OD_initial):
    save_path = os.path.dirname(os.path.realpath(__file__))
    pickle_path =  "%s/%s/%s.pickle" % (save_path,exp_name,exp_name)
    with open(pickle_path, 'w') as f:
        pickle.dump([start_time, OD_initial], f)

def graph_data(vials, exp_name, file_name):
    save_path = os.path.dirname(os.path.realpath(__file__))
    for x in vials:
        file_path =  "%s/%s/%s/vial%d_%s.txt" % (save_path,exp_name,file_name,x,file_name)
        data = np.genfromtxt(file_path, delimiter=',')
        plt.plot(data[:,0], data[:,1])
 ##      plt.ylim((-.05,.5))
        plot_path =  "%s/%s/%s_graph/vial%d_%s.png" % (save_path,exp_name,file_name,x,file_name)
        plt.savefig(plot_path)
        plt.clf()

def calc_growth_rate(vials, exp_name,elapsed_time, OD_maintain):
    save_path = os.path.dirname(os.path.realpath(__file__))
    for x in vials:
        ## Grab Data and make setpoint
        file_path =  "%s/%s/OD/vial%d_OD.txt" % (save_path,exp_name,x)
        OD_data = np.genfromtxt(file_path, delimiter=',')
        if np.shape(OD_data)[0] > 1010:
            time = OD_data[-1000:-1,0]
            raw_data = OD_data[-1000:-1,1]
            ## Smooth data
            window = 50
            mask = np.ones(window)/window
            raw_data = np.convolve(raw_data,mask, 'same')
            median_filter_range = 801;
            slope_data2 = movingslope(raw_data,10,1,10)
            slope_data2 = medfilt(slope_data2[:,0],median_filter_range);
            ## Calculate Average Growth for data
            average_data =((slope_data2[(median_filter_range/2):(np.size(slope_data2)-1-median_filter_range/2)])*3600)/OD_maintain
            ## Write Growth Rate to text
            log_path = "%s/%s/growth_rate/vial%d_growth_rate.txt" % (save_path,exp_name,x)
            text_file = open(log_path,"a+")
            text_file.write("%d,%s\n" %  (elapsed_time, np.average(average_data)))
            text_file.close()
        

def movingslope(vec,supportlength,modelorder,dt):
    n = np.size(vec);
    
    ##now build the filter coefficients to estimate the slope
    if ((supportlength % 2) == 1):
        parity = 1 #odd parity
    else:
        parity = 0;
    s = (supportlength-parity)/2
    t = np.arange((-s+1-parity),s+1)[np.newaxis]
    t = t.transpose()
    coef = getcoef(t,supportlength,modelorder)

    ## Apply the filter to the entire vector
    f = scipy.signal.lfilter(-coef,1,vec);
    Dvec = np.zeros((np.size(vec),1))
    r = s+ np.arange(1,(n-supportlength+2))
    for x in np.arange(0,np.size(r)):
        Dvec[r[x]] = f[supportlength+x-1]
        
    for i in np.arange(1,s+1):
        t = np.arange(1,supportlength+1)[np.newaxis]
        t = t - i
        t = t.transpose()
        coef = getcoef(t,supportlength,modelorder);
        coef = coef[np.newaxis]
        m = vec[0:supportlength][np.newaxis]
        m = np.transpose(m)
        Dvec[i-1]= np.dot(coef,m)

        if i<(s + parity):
            t = np.arange(1,supportlength+1)[np.newaxis]
            t = t - supportlength + i -1
            t = t.transpose()
            coef = getcoef(t,supportlength,modelorder)
            coef = coef[np.newaxis]
            m = vec[n-supportlength : n][np.newaxis]
            m = np.transpose(m)
            Dvec[n-i]= np.dot(coef,m)
            
    Dvec = Dvec/dt
    return Dvec


def getcoef(t,supportlength,modelorder):
    a = numpy.matlib.repmat(t,1,modelorder+1)
    b = numpy.matlib.repmat(np.arange(0,modelorder+1),supportlength,1)
    c = np.power(a,b)
    pinvA = numpy.linalg.pinv(c)
    coef = pinvA[1,:]
    return coef

def medfilt (x, k):
    """Apply a length-k median filter to a 1D array x.
    Boundaries are extended by repeating endpoints.
    """
    assert k % 2 == 1, "Median filter length must be odd."
    assert x.ndim == 1, "Input must be one-dimensional."
    k2 = (k - 1) // 2
    y = np.zeros ((len (x), k), dtype=x.dtype)
    y[:,k2] = x
    for i in range (k2):
        j = k2 - i
        y[j:,i] = x[:-j]
        y[:j,i] = x[0]
        y[:-j,-(i+1)] = x[j:]
        y[-j:,-(i+1)] = x[-1]
    return np.median (y, axis=1)

if __name__ == '__main__':
    temp_data = update_temp('23423423')
    print temp_data
```
