## Supplementary material for "Automated continuous evolution of proteins *in vivo*": Code associated with main paper: growthassayV3.m

```
function [DblTimeFinal,Lconf,Hconf] = growthassayV3(filename,r,c,r1,c1)
%Calculates the doubling time in hours of an excel output of a Tecan
% Magellan plate reader.
% 
% DblTimeFinal - doubling time in hours
% Lconf - 95% low bound confidence interval of doubling time
% Hconf - 95% high bound confidence interval of doubling time
%
% filename - excel input file
% r - number of rows measured in plate reader
% c - number of columns measured in pate reader
% r1 - number of rows to do analysis on starting from A1
% c1 - number of columns to do analysis on starting from A1

%% Part 1: Magellan import and organization of data
clearvars -except filename r c r1 c1
%Adjust file location, name, and number of time steps used for assay
fulldata = xlsread(filename,'Magellan Sheet 1');
rows = r;
cols = c;
tsteps = size(fulldata,1)/rows;


%Magellan time points
totaltime = tsteps*30-30;
tvector = linspace (0,totaltime,tsteps);

%Extract OD measurements into a 3D array (row,column,time)
for i = 1:rows
    for j = 1:cols
        for t = 1:tsteps
            k=i+rows*(t-1);
            OD(i,j,t) = fulldata(k,j);
        end
    end
end

%Normalize and log2 all data to minimum OD in well
for i = 1:rows
    for j = 1:cols
        for t = 1:tsteps
            ODnorm(i,j,t) = log2(OD(i,j,t)/min(squeeze(OD(i,j,:))));
        end
        
        growthA = reshape(ODnorm(i,j,:),1,length(ODnorm(i,j,:)));
        growthAs1 = smooth(growthA,'sgolay');
        growthAs2 = smooth(growthAs1,'moving');

        ODnorm(i,j,:) = growthAs2;
    end
end


%% Sliding window linear regression

%Constant sliding region with evolving fits, checking for highest slope and
%fit
ODnorm(isnan(ODnorm)) = 0; 
rows = r1;
cols = c1;

% Best growth rate in 5 hour interval
for totalPoints = 9:11
    for r = 1:rows
        for c = 1:cols
%             [r c]
            for slideStart = 1:(tsteps-totalPoints)
                currentFit = fit(transpose(tvector(slideStart:slideStart+totalPoints)),squeeze(ODnorm(r,c,slideStart:slideStart+totalPoints)),'poly1');
                coefficients = coeffvalues(currentFit);
                conf = confint(currentFit);
                
                mu = coefficients(1);
                err = (conf(2,1)-mu)/mu;            
                                
                
                if totalPoints == 9
                     muFit3(r,c,slideStart) = mu;
                     errFit3(r,c,slideStart) = err;
                elseif totalPoints == 10
                     muFit4(r,c,slideStart) = mu;
                     errFit4(r,c,slideStart) = err;
                elseif totalPoints == 11
                     muFit5(r,c,slideStart) = mu;
                     errFit5(r,c,slideStart) = err;
                end    
            end
        end
    end
end

%pick out highest mu and their errors
for r = 1:rows
    for c = 1:cols
        [mu3 index3] = max(muFit3(r,c,:));
        [mu4 index4] = max(muFit4(r,c,:));
        [mu5 index5] = max(muFit5(r,c,:));        
         muChoices(r,c,:) = [mu3 mu4 mu5];
         muChoicesStart(r,c,:) = [index3 index4 index5];
                
        Err3 = errFit3(r,c,index3);
        Err4 = errFit4(r,c,index4);
        Err5 = errFit5(r,c,index5);        
        ErrChoices(r,c,:) = [Err3,Err4,Err5];       
    end
end

%%
%Pick one mu for each well weighing both how high mu is and how low Error is. 
StdErr = (ErrChoices-muChoices)./muChoices;
for r = 1:rows
    for c = 1:cols
        if max(muChoices(r,c,:)) == muChoices(r,c,3) & min(ErrChoices(r,c,:)) == ErrChoices(r,c,3)
            muFinal(r,c) = muChoices(r,c,3);
            CritMatrix(r,c) = 3;
            plusorminus = ErrChoices(r,c,3);
            mulowBound(r,c)=muFinal(r,c)*(1-plusorminus);
            muhiBound(r,c)=muFinal(r,c)*(1+plusorminus);
        elseif max(muChoices(r,c,:)) == muChoices(r,c,2) & min(ErrChoices(r,c,:)) == ErrChoices(r,c,2)
            muFinal(r,c) = muChoices(r,c,2);
            CritMatrix(r,c) = 2;
            plusorminus = ErrChoices(r,c,2);
            mulowBound(r,c)=muFinal(r,c)*(1-plusorminus);
            muhiBound(r,c)=muFinal(r,c)*(1+plusorminus);
        elseif max(muChoices(r,c,:)) == muChoices(r,c,1) & min(ErrChoices(r,c,:)) == ErrChoices(r,c,1)
            muFinal(r,c) = muChoices(r,c,1);
            CritMatrix(r,c) = 1;
            plusorminus = ErrChoices(r,c,1);
            mulowBound(r,c)=muFinal(r,c)*(1-plusorminus);
            muhiBound(r,c)=muFinal(r,c)*(1+plusorminus);
        else
            [Crit iCrit] = min((ErrChoices(r,c,:)-muChoices(r,c,:))./(muChoices(r,c,:)));
            CritMatrix(r,c) = iCrit;
            muFinal(r,c) = muChoices(r,c,iCrit);
            mulowBound(r,c) = muFinal(r,c)*(1-ErrChoices(r,c,iCrit));
            muhiBound(r,c) = muFinal(r,c)*(1+ErrChoices(r,c,iCrit));
        end
        
    end
end

%return output variables
DblTimeFinal = 1./muFinal;
Lconf = 1./muhiBound; %reverse the bounds because 1/mu = dbltime
Hconf = 1./mulowBound; %reverse the bounds because 1/mu = dbltime

%% Plot and save


% Plot growth rates of all wells, highlighting linear region chosen
% Each well has separate subplot
fAll = figure(1);
for r = 1:rows
    for c = 1:cols
        subplot(8,12,(r-1)*12+c)
        pick = CritMatrix(r,c);
        Lcut = muChoicesStart(r,c,pick);
        Hcut = Lcut + (pick+5);
        plot(tvector(1:tsteps), squeeze(ODnorm(r,c,1:tsteps)));

        hold on;
        pick = CritMatrix(r,c);
        Lcut = muChoicesStart(r,c,pick);
        Hcut = Lcut + (pick+5);
        plot(tvector(Lcut:Hcut), squeeze(ODnorm(r,c,Lcut:Hcut)),'Color',[1 0 0],'LineWidth',1,'Marker','+');
        xlim([0 1500]);
        ylim([0 4]);
        hold off;
    end
end


figname1 = strcat(filename,'WholeRangePlot');
figname1PNG = strcat(figname1, '.png');

% Plot only linear region of growth rate for all wells on same graph
fLin = figure(2);
hold on
for r = 1:rows
    for c = 1:cols
        pick = CritMatrix(r,c);
        Lcut = muChoicesStart(r,c,pick);
        Hcut = Lcut + (pick+5);
        plot(tvector(Lcut:Hcut), squeeze(ODnorm(r,c,Lcut:Hcut)));
    end
end
hold off
figname2 = strcat(filename,'LinearRangePlot');
figname2PNG = strcat(figname2, '.png');


end
```
