## Supplementary material for "Automated continuous evolution of proteins *in vivo*": Code associated with main paper: main_eVOLVER.py

```
from Tkinter import *
from ttk import *
import eVOLVER_module
import time
import pickle
import os.path
import custom_script
import smtplib
import numpy as np


#Where the GUI is called and widgets are placed
class make_GUI:

    def __init__(self, master):
        master.wm_title("Liu Lab Experiments")
        note = Notebook(master)
        home = Frame(note)
        note.add(home, text = "Home")

        save_path = os.path.dirname(os.path.realpath(__file__))
        tabArray = [ ]
        
        for x in vials:
            newTab = Frame(note)
            note.add(newTab, text = "%d" % x)
            tabArray.append(newTab)
            
        self.quitButton = Button(home, text="Stop Measuring", command=stop_exp)
        self.quitButton.pack(side=BOTTOM)
        
        self.printButton = Button(home, text="Start/ Measure now", command=start_exp)
        self.printButton.pack(side=BOTTOM)

        note.pack(fill=BOTH, expand=YES)


## Task done by pressing printButton
def start_exp():
    stop_exp()
    update_Graphs()
    update_eVOLVER()

def stop_exp():
    global run_exp
    global graph_exp
    try:
        run_exp
    except NameError:
        print 
    else:
        print "Experiment Stopped!"
        root.after_cancel(run_exp)
        root.after_cancel(graph_exp)
    
## Updates Temperature and OD values (repeated every 10 seconds)
def update_eVOLVER():
    global OD_data, temp_data
    ##Read and record OD
    elapsed_time = round((time.time() - start_time)/3600,4)
    print "Time: %f Hours" % elapsed_time
    OD_data = eVOLVER_module.read_OD(vials)
    if OD_data == 'empty':
        print "Data Empty! Skipping data log..." 
    else:
        for x in vials: 
            OD_data[x] = OD_data[x] - OD_initial[x]
            if np.isnan(OD_data[x]):
                OD_data[x] = 1.5

    eVOLVER_module.parse_data(OD_data, elapsed_time,vials,exp_name, 'OD')

    ## Update and record temperature
    temp_data = eVOLVER_module.update_temp(vials,exp_name)
    eVOLVER_module.parse_data(temp_data, elapsed_time,vials,exp_name, 'temp')

    ##Make decision
    custom_functions(elapsed_time,exp_name)

    #Save Variables
    global run_exp
    eVOLVER_module.save_var(exp_name, start_time, OD_initial)
    run_exp = root.after(10000,update_eVOLVER)

### Update Graphs/ send alerts (repeats every 60 minutes)
def update_Graphs():
    elapsed_time = round((time.time() - start_time),0)
    # if OD_data == 'empty':
    #     send_alert()
  #  else:

  #      print "Increasing stir rate to prevent biofilm"
#         global graph_exp
# #        eVOLVER_module.graph_data(vials, exp_name, 'OD')
# #        eVOLVER_module.graph_data(vials, exp_name, 'temp')

#         MESSAGE = "20,20,20,20,20,20,20,20,20,20,20,20,20,20,20,20,"
#         eVOLVER_module.stir_rate(MESSAGE)
#         print "Fast stirring"
#         time.sleep(20)
#         MESSAGE = "0,0,0,0,0,0,0,0,0,0,0,0,0,0,0,0,"
#         eVOLVER_module.stir_rate(MESSAGE)
#         print "Normal stirring"
#         time.sleep(20)
#         graph_exp = root.after(60000*60,update_Graphs)
#         print "Clean cycle completed!"

#         eVOLVER_module.calc_growth_rate(vials, exp_name,elapsed_time,.3)
#         MESSAGE = "0,0,0,0,0,0,0,0,0,0,0,0,0,0,0,0," 
#         eVOLVER_module.stir_rate(MESSAGE)

#         time.sleep(10)

#         print "Efflux ON for overflow check"
#         MESSAGE =  "111111110000000000000000,0,10,"
#         eVOLVER_module.fluid_command(MESSAGE, 0, elapsed_time, 1, exp_name, 1,'n')
#         MESSAGE =  "11111111000000000000000000000000,0,10,"
#         eVOLVER_module.fluid_command(MESSAGE, 0, elapsed_time, 1, exp_name, 1,'n')

#         time.sleep(60)

    

#### Custom Script
def custom_functions(elapsed_time, exp_name):
    global OD_data, temp_data
    if OD_data == 'empty':
        print "UDP Empty, did not execute program!"
    else:
        ###load script from another python file
        custom_script.test(OD_data, temp_data, vials,elapsed_time, exp_name)
    
    


### Runs if this is main script 
if __name__ == '__main__':
    global graph_exp
    exp_name = 'ZZ_expt_MST_PID_V4_HisA_2018_08_30_RE'
    vials = range(0,16)
    start_time, OD_initial  =  eVOLVER_module.initialize_exp(exp_name,vials)
    root=Tk()
    make_GUI(root)
    graph_exp = root.after(60000*5,update_Graphs)
    update_eVOLVER()
    root.mainloop()
```
